# FIDDLE: a deep learning method for chemical formulas prediction from tandem mass spectra

**DOI:** 10.1101/2024.11.25.625316

**Authors:** Yuhui Hong, Sujun Li, Yuzhen Ye, Haixu Tang

**Affiliations:** Luddy School of Informatics, Computing, and Engineering, Indiana University Bloomington, 700 N. Woodlawn Avenue, Bloomington, 47408, Indiana, USA

## Abstract

Molecular identification through tandem mass spectrometry is fundamental in small molecule analysis, with formula identification serving as an initial step in the process. Current computational methods often struggle with accuracy, speed, and scalability for relatively larger molecules, limiting high-throughput workflows. We present FIDDLE (**F**ormula **ID**entification by **D**eep **LE**arning), a deep learning-based method trained on over 38,000 molecules and 1 million MS/MS spectra from various Quadrupole Time-of-Flight (Q-TOF) and Orbitrap instruments. FIDDLE accelerates formula identification by more than 10-fold and achieves top-1 and top-5 accuracies of 88.3% and 93.6%, respectively, outperforming state-of-the-art methods based on top-down (SIRIUS) and bottom-up (BUDDY) approaches by over 10%. On external metabolomics datasets, FIDDLE achieves top-5 accuracies of 75.1% (positive ion mode) and 66.2% (negative ion mode), with further improvements to 80.0% and 73.8% when combined with SIRIUS and BUDDY.

## 1 Introduction

Tandem mass spectrometry (MS/MS) is an essential analytical tool for identifying small molecules and elucidating their structural characteristics. The standard approach for identifying unknown analytes from MS/MS data involves searching spectra against reference spectral libraries [1–3]. However, due to limitations in time, labor, and resources, a significant portion of chemical signatures remains uncharacterized, often termed ‘dark matter’ [4]. These unidentified small molecules may possess unique bioactivities and play crucial roles in understanding biological mechanisms. Unfortunately, such molecules may lack corresponding reference spectra in spectral libraries or may not have been previously reported in the literature (i.e., the ‘unknown unknowns’ [5]). As a result, the identification of unknown compounds has become a challenging yet vital research area, spanning metabolomics, environmental analysis, natural product and drug discovery, and more [6–11]. Specifically, the prediction of molecular formulas serves as the initial and most fundamental step, providing critical constraints that facilitate structural elucidation and the annotation of fragments for these unknown compounds [12, 13].

While MZmine initially incorporated isotope pattern matching, MS/MS fragmentation analysis, and heuristic rules as a toolbox [14], computational methodologies for chemical formula identification from MS/MS data have evolved into top-down and bottom-up approaches, exemplified by SIRIUS [15] and BUDDY [16], respectively. SIRIUS begins by generating candidate formulas through the analysis of isotope patterns [17], then computes a fragmentation tree for each candidate [18] and evaluates them by comparing theoretical fragmentation patterns against experimental MS/MS data. The evaluation metric considers multiple factors including fragment masses, intensities, and isotope patterns to estimate the likelihood of each candidate producing the observed spectrum. However, SIRIUS’s reliance on neutral loss fragments limits its ability to handle multiply-charged spectra (7.7% and 1.8% in preprocessed NIST23 Q-TOF and Orbitrap datasets, as shown in Supplementary Fig. 1), which contain charged loss fragments. Additionally, its efficiency is hindered by the computational demand of generating fragmentation trees for all potential formulas inferred from isotope patterns. MIST-CF shows that fragmentation trees can be replaced with a simple peak subformula assignment routine, achieving equally accurate and fast predictions [19]. However, it still relies on SIRIUS’s algorithmic decomposition of exact masses into formula candidates limiting efficiency and accuracy. In contrast, BUDDY significantly reduces the number of candidate formulas by focusing on those explainable by MS/MS data, using a reference library of known formulas. It ranks candidates matching the precursor mass and estimates a false discovery rate to provide a confidence score. However, BUDDY’s scope is restricted by the coverage of its reference library, potentially missing entirely uncharacterized and previously unreported formulas. Our analysis illustrated in Supplementary Fig. 2 revealed that 45 unique formulas represented in MS/MS spectra in the National Institute of Standards and Technology (NIST) Spectral Library, Massbank of North America (MoNA) [2], Global Natural Product Social Molecular Networking (GNPS) Spectral Library [20], and the Agilent Personal Compound Database and Library (PCDL) fall outside the MS/MS-explainable space of BUDDY, rendering them unanalyzable by the method.

Both computational methods underutilize the full scope of information present in MS/MS spectrum data. SIRIUS 6, for instance, considers a limited number of peaks (up to 60), while BUDDY relies on manually extracted features, such as double-bond equivalent values of annotated fragments. As a result, increasing precursor mass-to-charge ratio (*m/z*) leads to higher computational complexity and significantly decreased accuracy due to the exponentially growing number of candidate molecular formulas, which expands the search space and increases ambiguity. For instance, at *m/z* 800, the number of candidates reaches tens for BUDDY and tens of thousands for SIRIUS [16]. This limitation stems from the fact that higher precursor *m/z* values often correspond to larger, more complex molecular structures, when then requires the evaluation of a broader range of potential formulas. Furthermore, the peaks excluded from analysis, along with the relationships between considered and unconsidered peaks, may hold crucial structural information that is not exploited.

In this paper, we address these limitations by introducing a deep learning approach to chemical formula identification. We present FIDDLE (**F**ormula **ID**entification from tandem mass spectra by **D**eep **LE**arning), which employs dilated convolutions with large kernels [21, 22] to extract high-dimensional representations of MS/MS data using extremely large receptive fields. To predict candidate formulas, the model is trained using a composite objective that includes a primary formula regression loss, a contrastive loss, and auxiliary task losses to enhance performance. These initial predictions are refined using a breadth-first search algorithm that adjusts atomic compositions to align the candidate formulas with the precursor mass. Additionally, we train a secondary deep learning model to estimate confidence scores for candidate formulas and rank them based on MS/MS features learned from the formula identification model. Compared to traditional computational methods, our approach dramatically reduces the candidate formula space for a given MS/MS spectrum to a small number (at most 5 formulas by default). This reduction is facilitated by the deep learning model, which benefits from accelerated GPU-based tensor computations. The narrowed candidate space simplifies confidence score estimation, and the MS/MS features learned by the deep learning model can be reused for this purpose.

## 2 Results

### 2.1 Deep learning method for formula identification

Predicting target formulas directly from MS/MS spectra under varying experimental conditions presents significant challenges. To address this, we break down the task into three steps as illustrated in Fig. 1 (a): 1) predicting formulas from MS/MS spectra using a deep learning model; 2) generating candidate formulas using a breadth-first formula refinement algorithm; and 3) calculating confidence scores for the candidate formulas using an additional deep learning model. The formula refinement step relaxes the requirement for exceptionally high initial prediction accuracy by allowing adjustments to candidate formulas with minimal atom modifications. Moreover, since assessing the correctness of a limited set of predictions is easier than identifying the top correct outcomes from an infinitely large pool [23], the refinement step also improves the confidence score estimation.

**Fig. 1:**
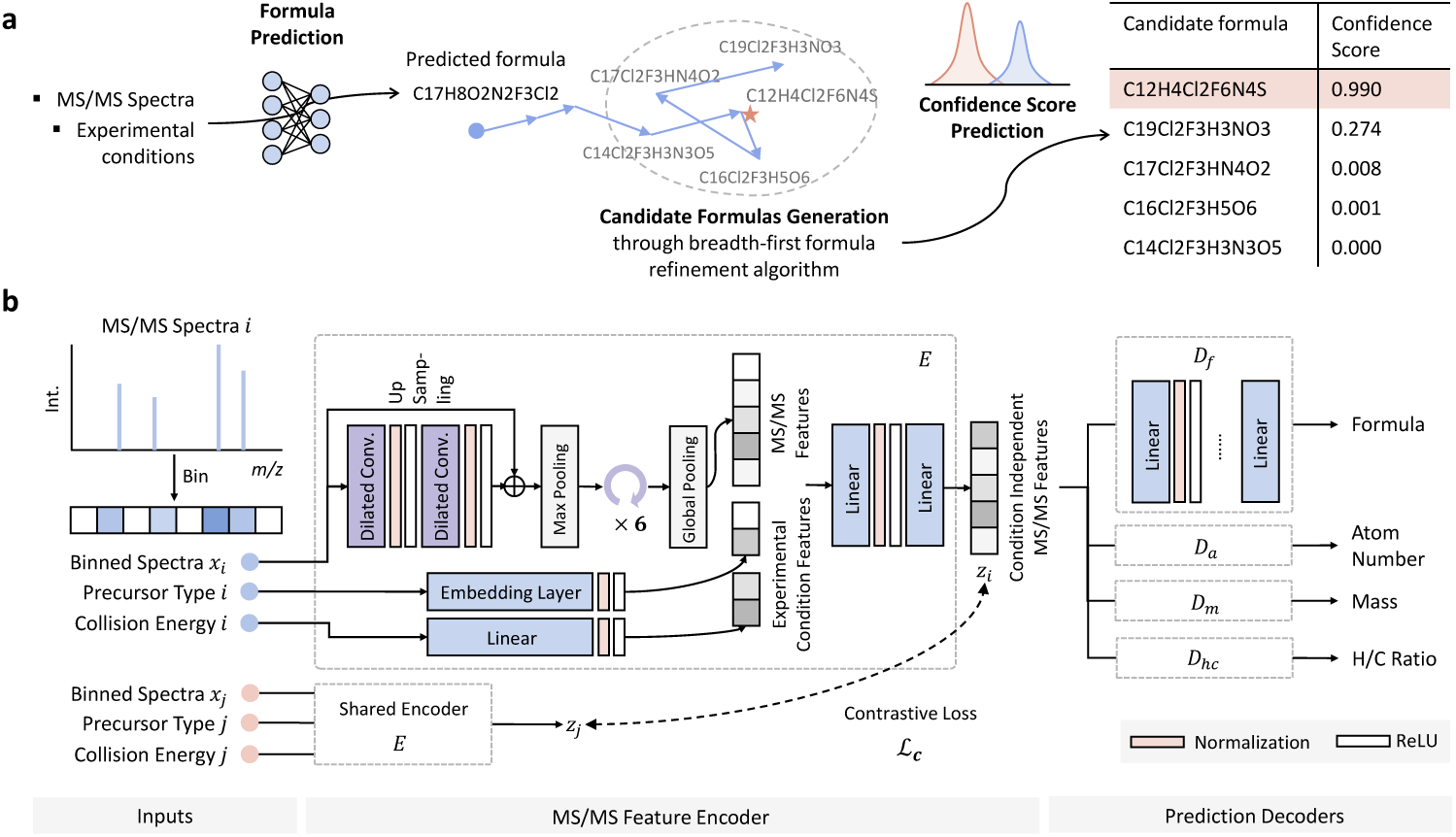
formula identification from tandem mass spectra using deep learning. (a) The FIDDLE workflow comprises three main steps: predicting formulas from MS/MS spectra using a deep learning model; generating candidate formulas through a breadth-first formula refinement algorithm; and predicting confidence scores for the candidate formulas. (b) The deep learning model architecture includes an MS/MS spectrum encoder (*E*) and decoders (*D_f_*, *D_a_*, *D_m_*, and *D_hc_*) that outputs the predicted formula along with auxiliary variables, such as atom count, molecular mass, and H/C ratio.

To input an MS/MS spectrum into a deep learning model, we first bin it into a 1-D vector with a fixed mass-to-charge ratio (*m/z*) resolution. For example, an MS/MS spectrum with a maximum *m/z* of 1500 Da is binned into a vector of length 7500, with each bin representing a resolution of 0.2 Da. Molecular formulas are directly converted into formula vectors, where each element type is represented by its atom count as the corresponding value in the vector. For instance, the molecular formula C_6_H_12_O_6_ can be represented as the vector [6, 12, 6, 0*, …*], where the first three integers correspond to the number of carbon, hydrogen, and oxygen atoms, respectively, followed by atom counts for other elements in a predefined order.

To encode the MS/MS spectra, we use stacked dilated convolutions with large kernels to capture relationships between peaks across broad mass ranges [21, 22]. This technique expands the model’s receptive field, enabling it to analyze local and global spectral patterns at the same time. It serves as a powerful and computationally efficient alternative to fully connected layers. The learned MS/MS features are then concatenated with experimental conditions, such as collision energies, precursor types, and experimental precursor *m/z*, and fed into two linear layers to produce condition-independent MS/MS features, denoted as **z_i_** and **z_j_** in Fig. 1 (b). During training, a contrastive loss is applied to ensure that these condition-independent MS/MS features are close for spectra from the same molecule and far apart for spectra from different molecules. We use sequential linear layers as decoders for both formula identification and auxiliary tasks, including atom number prediction, molecular mass prediction, and H/C ratio prediction. These auxiliary tasks, incorporated through multitask learning, enhance the generalizability of the deep learning model and serve as a form of regularization [24]. The detailed architecture and parameters of the model are specified in Section 4.2.

Recognizing that deep learning models cannot guarantee the validity or perfect accuracy of predicted formulas, we developed a breadth-first formula refinement algorithm. This algorithm aims to make minimal adjustments to atom counts to ensure that the formulas comply with SENIOR rules [25] and align with the target mass within a specified mass tolerance—specifically, 10 parts per million (ppm) for MS/MS data from Q-TOF instruments and 5 ppm for data from Orbitrap instruments. This refinement process produces a set of *k* candidate formulas for each MS/MS spectrum, where *k* is set to 5 by default. Note that the algorithm is flexible enough to integrate results from various formula identification methods, such as SIRIUS and BUDDY, where the predicted formula can be expanded into a longer list of candidate formulas. An auxiliary model is then developed to estimate confidence scores using the MS/MS features learned during formula identification and each candidate formula (see details in Section 4.4). Finally, the candidate formulas are ranked based on their estimated confidence scores.

### 2.2 Performance of formula identification

MS/MS spectra were collected from NIST23, NIST20, Agilent PCDL, MoNA, and GNPS, as well as from an internal dataset (see Section 4.1.1 for details) acquired using a Waters Q-TOF mass spectrometer. The MS/MS spectra were preprocessed according to the methods described in Section 4.1, including filtering based on peak count, molecular mass, atom type and number, and mass difference in ppm. Additional preprocessing steps included simulating precursor *m/z* values for the NIST dataset, simplifying precursor types, and constructing the training set for contrastive learning. In total, 131,224 MS/MS spectra from 15,399 molecules acquired with Q-TOF mass spectrometers and 965,656 MS/MS spectra from 28,383 molecules acquired with Orbitrap mass spectrometers were used for training and evaluation. A summary of the number of spectra and compounds in each dataset is provided in Table 1. We retained MS/MS spectra from compounds (represented as canonical SMILES without stereochemical information) found exclusively in NIST23 and not in any other libraries (including NIST20) as the test set, ensuring these spectra were not used during the training of any of the models being compared. It is important to note that these spectra were published after the release of BUDDY and SIRIUS. Hence, it is important not to leverage them to train FIDDLE and maintain a fair comparison. Consistent with previous studies [16], we used the *top K accuracy* to evaluate the performance of formula identification algorithms. This metric is calculated as the proportion of spectra for which the correct formula is included among the top *K* (by default *K* = 5) ranked formulas predicted by a given algorithm. The settings for comparison methods are specified in Section 4.6.

**Table 1:**
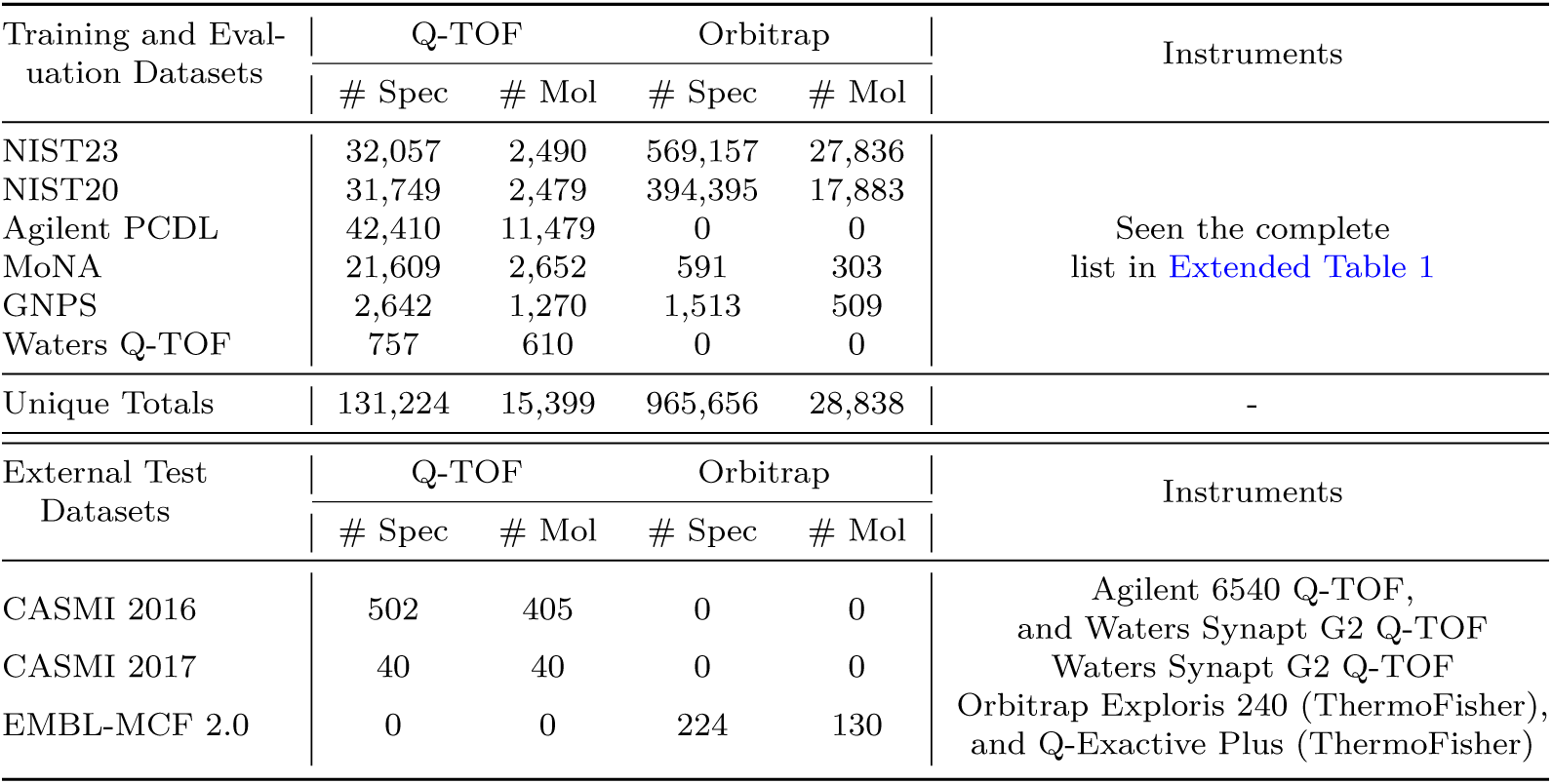
Total numbers of spectra and compounds used for training, evaluation, and external test.

As shown in Fig. 2 (a)-(f), FIDDLE outperforms the other state-of-the-art formula identification algorithms (BUDDY and SIRIUS), and speeds up the cumulative runtime by approximately 10-fold compared to BUDDY and 100-fold compared to SIRIUS. In addition to formula identification, performance metrics for auxiliary tasks, including mass, atom number, and H/C ratio predictions, are presented in Extended Fig. 1. Notably, the top-1 accuracy of BUDDY and SIRIUS declines significantly for larger compounds (molecular weight *>* 800 Da; Fig. 2 (b) and (e)), with BUDDY’s accuracy decreasing to 0.427 for Q-TOF and 0.684 for Orbitrap, and SIRIUS’s accuracy decreasing to 0.187 for Q-TOF and no output within the timeout limit for Orbitrap, compared to 0.844, 0.702, 0.583, and 0.669, respectively on smaller compounds (Fig. 2 (a) and (d)). In contrast, FIDDLE maintains robust performance on large compounds, achieving top-1 accuracies of 0.642 and 0.813 for Q-TOF and Orbitrap spectra, respectively. Notably, for small compounds, FIDDLE slightly outperforms BUDDY and SIRIUS, achieving top-1 accuracies of 0.906 and 0.881 for Q-TOF and Orbitrap, respectively. In top-5 formula prediction, FIDDLE also consistently outperformed BUDDY and SIRIUS, especially on challenging large compounds. For Q-TOF data, FIDDLE’s accuracy on large compounds (0.754) was substantially higher than that of BUDDY (0.489) and SIRIUS (0.178). This trend was even more substantial for Orbitrap data, where FIDDLE achieved an accuracy of 0.971 for large compounds, while BUDDY scored 0.684 and SIRIUS failed to return a result in time. Moreover, incorporating BUDDY’s results into FIDDLE’s candidate formula pool further improved prediction accuracy, indicating that BUDDY and FIDDLE perform well on different sets of compounds and can be complementary. While combining these methods may require additional running time, it can yield superior overall results. The best accuracy and running time of FIDDLE for different settings of K on Q-TOF test spectra are shown in Fig. 2 (h). Higher *K* values improve accuracy at the cost of increased computational time, enabling users to balance performance and efficiency based on their requirements.

**Fig. 2:**
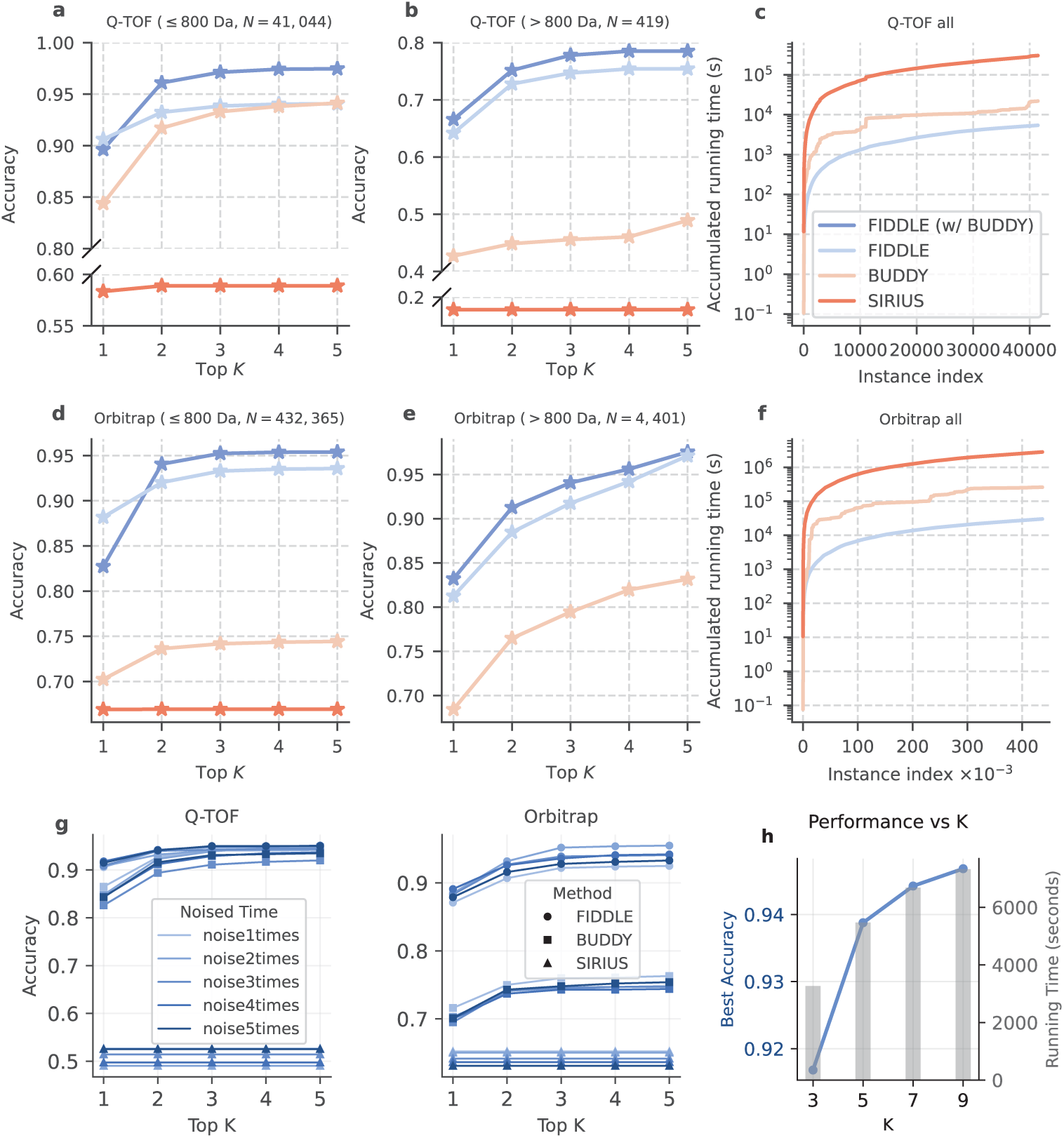
Comparison of the performance of FIDDLE with SIRIUS and BUDDY on formula identification. (a-b) Top *K* accuracy for formula identification on Q-TOF MS/MS spectra of molecules ≤ 800 Da and *>* 800 Da, respectively. *N* represents the number of spectra. (c-f) Same analysis for Orbitrap spectra. SIRIUS produced no output for large Orbitrap molecules due to runtime limits. FIDDLE supports all 14 precursor types (charges +1, -1, +2), while BUDDY and SIRIUS are evaluated only on supported types, excluding timeouts. (g) Top *K* accuracy on the MS/MS spectra with different levels of added noise. (h) FIDDLE’s performance across different *K* settings, where the left y-axis (blue lines) represents the best accuracy of top *K* candidates and the right y-axis (gray bars) represents accumulated running times.

We assessed FIDDLE’s generalizability by comparing its performance on two distinct training and test set splits: one divided randomly by canonical SMILES and a more stringent split divided by unique chemical formula. The formula-based split is significantly more challenging, as it requires the model to predict formulas it has never encountered during training. Despite this, FIDDLE’s accuracy decreased by only about 10% on this task compared to the standard SMILES-based split (*p <* 0.001, one-sample t-test), demonstrating robust performance. Detailed results are shown in Supplementary Fig. 3.

To assess robustness to spectral noise, we evaluated FIDDLE, BUDDY, and SIRIUS on 1,000 randomly sampled MS/MS spectra from the test set. We systematically added Gaussian noise at five increasing levels, with detailed methods described in Section 4.1.4. FIDDLE demonstrated superior noise resilience across both Q-TOF and Orbitrap instruments, maintaining over 90% accuracy even under large noise conditions. On Q-TOF spectra, FIDDLE achieves 95.0% top-5 accuracy at the highest noise level compared to BUDDY’s 93.5% and SIRIUS’s 52.5%. On Orbitrap spectra, FIDDLE maintains 93.3% accuracy while BUDDY decreases to 75.4% and SIRIUS to 63.1%. Overall, FIDDLE exhibited minimal performance degradation as noise increased, whereas BUDDY showed a moderate decline and SIRIUS performed poorly but consistently, underscoring FIDDLE’s practical advantage in real-world scenarios with varying spectral quality.

Additional evaluation on chimeric spectra—synthetic mixtures of authentic MS/MS data [26]—demonstrates the expected trade-off between FIDDLE’s predictive performance and robustness to signal mixtures (Supplementary Note 1). These results inform potential improvements through data augmentation strategies.

It is worth noting that interpretability can be a limitation of deep learning models compared to computation-based methods; therefore, we provide a potential interpretation method for FIDDLE in Supplementary Note 4 using SHAP [27]. However, current interpretation approaches face constraints from binned spectral resolution and limitations in providing specific atom counts for direct fragment annotation. Future extensions to molecular structure prediction could offer more intuitive explanations by directly linking spectral features to structural fragments.

### 2.3 Impact of data characteristics on model performance

We conducted a comprehensive analysis to evaluate how model performance is influenced by various data characteristics. The factors we investigated include experimental metadata (such as collision energy and precursor type), molecular properties (like size and chemical class), and the internal MS/MS representations learned by FIDDLE.

#### Metadata

We integrated precursor type and collision energy as metadata in the deep learning model and evaluated performance under different conditions, as shown in Fig. 3 (a) and (b). Performance exhibited a positive correlation with training dataset size, with the following examples illustrating this relationship. FIDDLE demonstrates optimal performance when analyzing spectra of dimers [2 M + H]^+^ and [2 M – H]^-^, perhaps due to their more predictable fragmentation patterns and simpler spectral signatures (Extended Fig. 2). Furthermore, higher collision energies produce more peaks, enhancing pattern recognition and boosting performance (the circle sizes denote the peak numbers). Only the subset of spectra with collision energies in the range [40, ∞) for the Q-TOF instrument had sufficient training data for optimal results by FIDDLE.

**Fig. 3:**
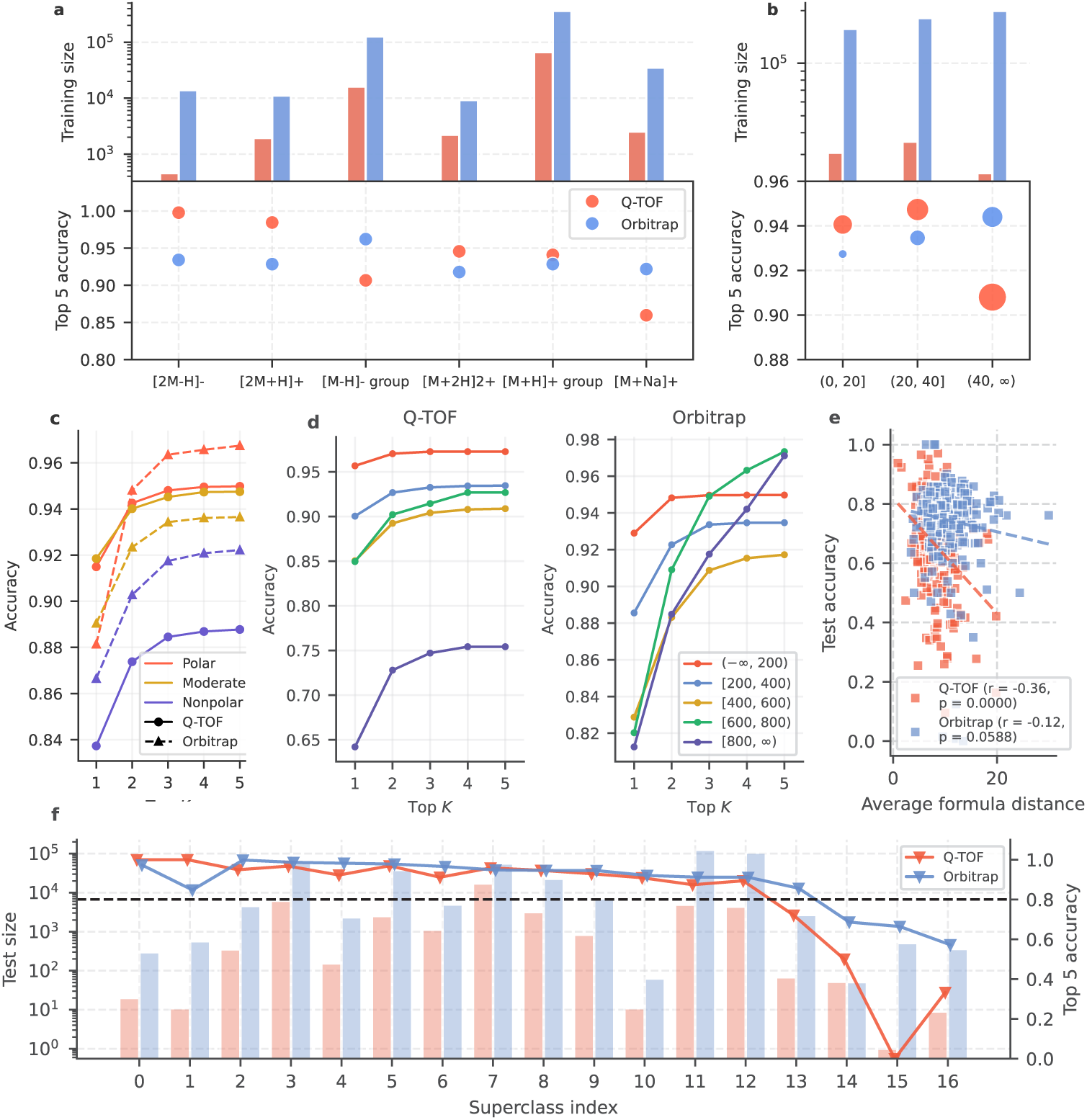
Performance across metadata (a and b), molecular (c, d, and f), and MS/MS representation (e) factors. (a,b) Top-5 accuracy by precursor type and collision energy. (c,d) Top-5 accuracy by molecular polarity (LogP) and mass range. (e) Accuracy vs. formula distance for similar MS/MS spectra. (f) Top-5 accuracy across 17 chemical superclasses (indices in Supplementary Note 2).

#### Molecular polarity

LogP values were computed from SMILES strings using RDKit’s Crippen.MolLogP method. Compounds were classified into three polarity groups: Polar (LogP *<* 0), Moderate (0 ≤ LogP *<* 3), and Nonpolar (LogP ≥ 3). Identification accuracy exhibited a strong inverse correlation with LogP values (Fig. 3 c). Polar compounds achieved the highest Top-5 accuracy (95.0%), followed by moderate (94.7%) and nonpolar compounds (88.8%), representing a 6.2% performance gap that was consistent across both Q-TOF and Orbitrap instruments. This trend is likely due to the superior ionization efficiency and more consistent fragmentation that polar compounds exhibit under electrospray ionization (ESI) mass spectrometry.

#### Molecular mass

Molecular weights were calculated from SMILES using RDKit’s MolWt function and classified into five groups: *<* 200, 200-400, 400-600, 600-800, and ≥ 800 Da (Fig. 3 d). Accuracy exhibited a complex relationship with molecular mass, with the smallest compounds (*<* 200 Da) achieving the highest performance (97.3% Top-5 accuracy for Q-TOF, 95.0% for Orbitrap). While the model performance on Q-TOF data declined substantially for larger molecules (≥ 800 Da: 75.4%), on Orbitrap data, the model maintained robust accuracy across all mass ranges, performing particularly well for the largest compounds (≥ 800 Da: 97.1%). This performance divergence suggests that the Orbitrap’s superior mass resolution is key to identifying large molecules, whose richer fragmentation patterns provide discriminative features that deep learning methods can effectively exploit.

#### Molecular types

Test compounds were classified using ClassyFire Batch into 17 superclasses (see Supplementary Note 2). Fig. 3 (f) reveals substantial variation in identification accuracy across chemical superclasses. While most superclasses achieved high performance (>90% Top-5 accuracy), four categories showed notably lower performance: Organosulfur compounds (72.1% on Q-TOF data and 85.6% on Orbitrap data), Organic Polymers (50.0% on Q-TOF data and 68.6% on Orbitrap data), Organohalogen compounds (0% on Q-TOF data and 66.4% on Orbitrap data), and Unknown compounds (33.3% on Q-TOF data and 57.2% on Orbitrap data). This drop in performance is likely due to insufficient training data for these underrepresented superclasses.

#### MS/MS representation space

To visualize the learned feature space, we applied t-SNE to project FIDDLE’s 512-dimensional latent vectors into a 2D representation, which we then divided into a 20×20 grid. For each cell in the grid, we calculated the top-1 prediction accuracy and the average Euclidean distance between molecular formulas. For cells containing more than 1,000 formula pairs, we used a random sample of 1,000 for this calculation. As shown in Fig. 3 (e), accuracy is negatively correlated with average formula distance. This correlation was highly significant for Q-TOF data (Pearson’s *p* « 0.0001), but not statistically significant for Orbitrap data (*p* = 0.0588). This finding confirms that FIDDLE’s prediction accuracy is challenged in regions where similar spectral representations map to divergent formulas, indicating that the learned features in these cases are not distinctive enough for discrimination. A detailed analysis is provided in Supplementary Note 3.

### 2.4 Ablation study of FIDDLE’s components

The MoNA (Q-TOF) dataset is used to illustrate the improvements of FIDDLE achieved through data augmentation, contrastive loss, and post-processing steps, including candidate formula generation and confidence score prediction. We evaluated FIDDLE’s performance across different processing steps by measuring the proportion of top-1 formulas with varying numbers of missed atoms (Fig. 4 (a)) and missed heavy atoms (Fig. 4 (c)), respectively. Because hydrogen (approximately 1.008 Da) is light and difficult to accurately determine, the count of heavy atoms (excluding hydrogen) is also considered. The number of missed atoms is calculated by summing the differences in atom counts across all elements (e.g., carbon, oxygen, nitrogen). A comparison of FIDDLE’s performance in ‘Pred w/o DA’ (prediction without data augmentation) and ‘Pred’ (prediction with data augmentation) shows that data augmentation significantly increases the proportion of correctly predicted formulas (from 0% to 9% for all atoms, and from 18% to 26% for heavy atoms). The contrastive loss further enhances performance, as seen in results labeled ‘Pred w/o CL’. Comparing the performance of FIDDLE under ‘Pred’ with ‘Post Top-1’ (top-1 accuracy after post-processing) to ‘Post Top-5’ (top-5 accuracy after post-processing) shows that FIDDLE’s candidate formulas cover the correct formulas for more than 84% of the MS/MS spectra. These candidates are subsequently ranked based on their confidence scores predicted by FIDDLE, achieving an AUC (area under the ROC curve) of 0.97, as shown in Fig. 4 (b). From Fig. 4 (d), it is clear that correct and incorrect formulas can be effectively distinguished based on their confidence scores. Notably, after post-processing, no formulas with three or fewer missed atoms remain, indicating that while post-processing may occasionally worsen certain incorrect predictions, most incorrect formulas with only a small number of missing atoms are effectively refined.

**Fig. 4:**
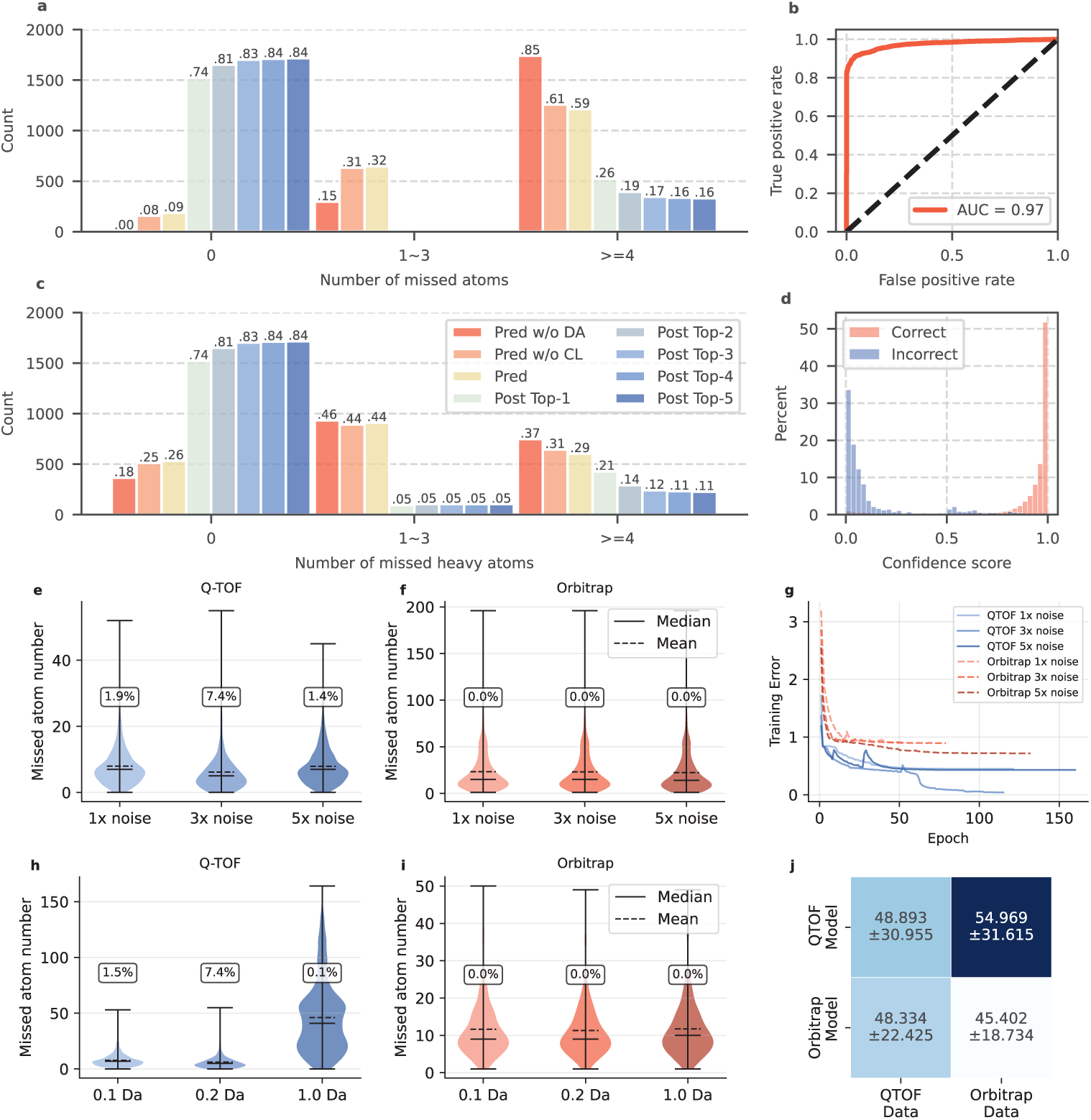
Ablation study of data augmentation, contrastive loss, post-processing, and hyperparameter in FIDDLE. (a) and (c) show the proportion of top-1 formulas with different numbers of missed atoms (a) and missed heavy atoms (c), respectively, predicted by FIDDLE with different processing steps, where ‘Pred w/o DA’ indicates the model trained without data augmentation; ‘Pred w/o CL’ indicates the model trained without contrastive learning; ‘Pred’ indicates the model trained with contrastive loss; ‘Post Top-1’ to ‘Post Top-4’ indicate the top-1 to top-4 formulas from post-processing. (b) and (d) show the AUC-ROC curve (b) and the distributions (d) for discriminating correctly and incorrectly predicted formulas based on their predicted confidence scores among the top-5 predicted formulas from the MS/MS spectra in the MoNA dataset. (e) and (f) show the performance of the models trained on MS/MS spectra with different levels of added noise (formula-level accuracy is annotated in the rounded rect), while (g) shows their corresponding learning curves. (h) and (i) show the performance of the model trained with different resolutions. (j) shows cross-instrument validation (missed atom number) on identical compound sets (diagonal: train and test on the same instrument type; off-diagonal: train and test on different instrument types).

We experimentally optimized the noise augmentation parameters of FIDDLE by training formula prediction models with different noise intensities applied to the training spectra (1×, 3×, and 5× multipliers). The MoNA (QTOF) dataset is reused in these experiments, and 20,000 spectra from NIST23 (Orbitrap) are randomly selected and split using the strategy described in Section 4.1.4. The results reveal an instrument-dependent response: for Q-TOF data, moderate noise augmentation (3×) yielded the best performance, whereas both lower (1×) and higher (5×) noise levels led to suboptimal model convergence. In contrast, Orbitrap spectra did not benefit from noise augmentation at any tested level, suggesting that adding Gaussian noise may compromise the inherent spectral consistency of high-resolution instruments (Extended Fig. 4). The resolution of input spectra is also experimentally optimized as shown in Fig. 4 (h) and (i), where 0.2 Da resolution achieves the best performance on both QTOF and Orbitrap instruments. Orbitrap shows robustness across different resolutions, with the largest number of missed atoms occurring at a resolution of 1 Da.

To investigate the effect of instrument type, we construct subsets from MoNA (QTOF) and NIST23 (Orbitrap), where the spectra are from the same 2357 compounds but acquired on different instruments. The spectra from Orbitrap are downsampled to 18,830, the same size as the QTOF spectra. The split strategy is described in Section 4.1.4. Then we conduct the same-instrument validation and cross-instrument validation (Fig. 4 j). Interestingly, the Orbitrap model performs better on both QTOF and Orbitrap data, likely because it is trained on higher-resolution spectra with more detailed spectral features, enabling it to learn more robust representations that generalize across platforms. While this demonstrates the advantages of high-resolution training data, QTOF-specific models remain valuable for laboratories with limited access to Orbitrap instruments and for high-throughput applications where faster acquisition is preferred.

### 2.5 Evaluation using benchmarking metabolite datasets

Next, we compared the performance of FIDDLE against SIRUIS, BUDDY, and MISTCF on three benchmarking metabolite datasets, the Critical Assessment of Small Molecule Identification (CASMI) 2016 [28], CASMI 2017, and EMBL-MCF 2.0 [29] datasets, respectively. For a fair comparison, we removed overlapping compounds in these datasets with our training set; in total 181 (231 spectra), 2 (2 spectra), and 107 (184 spectra) were removed from these three datasets, respectively (for details see Table 1). For a fair comparison, we train MIST-CF* using the same training and test data as used for FIDDLE*, excluding unsupported precursor types from both datasets (see details in Extended Table 4).

As illustrated in Fig. 5, FIDDLE performs comparably well with SIRIUS and BUDDY on the two CASMI datasets, while demonstrating significantly better performance on the EMBL-MCF 2.0 dataset. Since all methods achieve near-optimal performance on top-3 accuracy, with only FIDDLE (w/ BUDDY & SIRIUS), BUDDY, and MIST-CF* showing less than 2% improvement from top-5 to top-3, top-4 and top-5 accuracies are not demonstrated here. Complete top-5 accuracy results are provided in Supplementary Fig. 4 and 5. The slightly worse performance of FIDDLE on CASMI can be attributed to multiple factors, including different data acquisition protocols, lower MS/MS spectral quality (CASMI 2017 used Q-TOF, while CASMI 2016 and EMBL used Orbitrap), lower compound similarity to the training set (see Extended Fig. 3 (a)), spectral complexity, among others. The computation-based methods exhibit greater sensitivity to mass deviation, as their performance decreases on the EMBL-MCF 2.0 dataset with larger mass deviations compared to the CASMI dataset, as shown in Extended Fig. 3 (b). Notably, incorporating the NIST23 dataset into the training set enhances FIDDLE’s performance on these external test sets primarily due to the larger training data volume (see analysis in Supplementary Note 4), resulting in higher or comparable top-3 accuracies compared to SIRIUS, BUDDY, and MIST-CF. As shown in Fig. 5 (g-i), FIDDLE and BUDDY each performs better at different formulas and exhibit distinct error patterns (e.g., C_4_S and O_5_ for BUDDY versus C_3_ and HOF for FIDDLE); therefore, combining candidate formulas from all methods and ranking them by FIDDLE’s predicted confidence scores further improves performance. The challenging component pairs for both BUDDY and FIDDLE, such as N_3_ and C_2_H_2_O (which appear as N_6_ and C_4_H_4_O_2_ in BUDDY’s error patterns) are worth further investigations. On average, across three test sets, our method achieves top-5 accuracies of 80.0% (positive) and 73.8% (negative) ion mode, while BUDDY, SIRIUS, and MIST-CF achieve average top-5 accuracies of 69.9% (positive) and 61.4% (negative), 69.3% (positive) and 67.6% (negative), and 66.5% (positive), respectively.

**Fig. 5:**
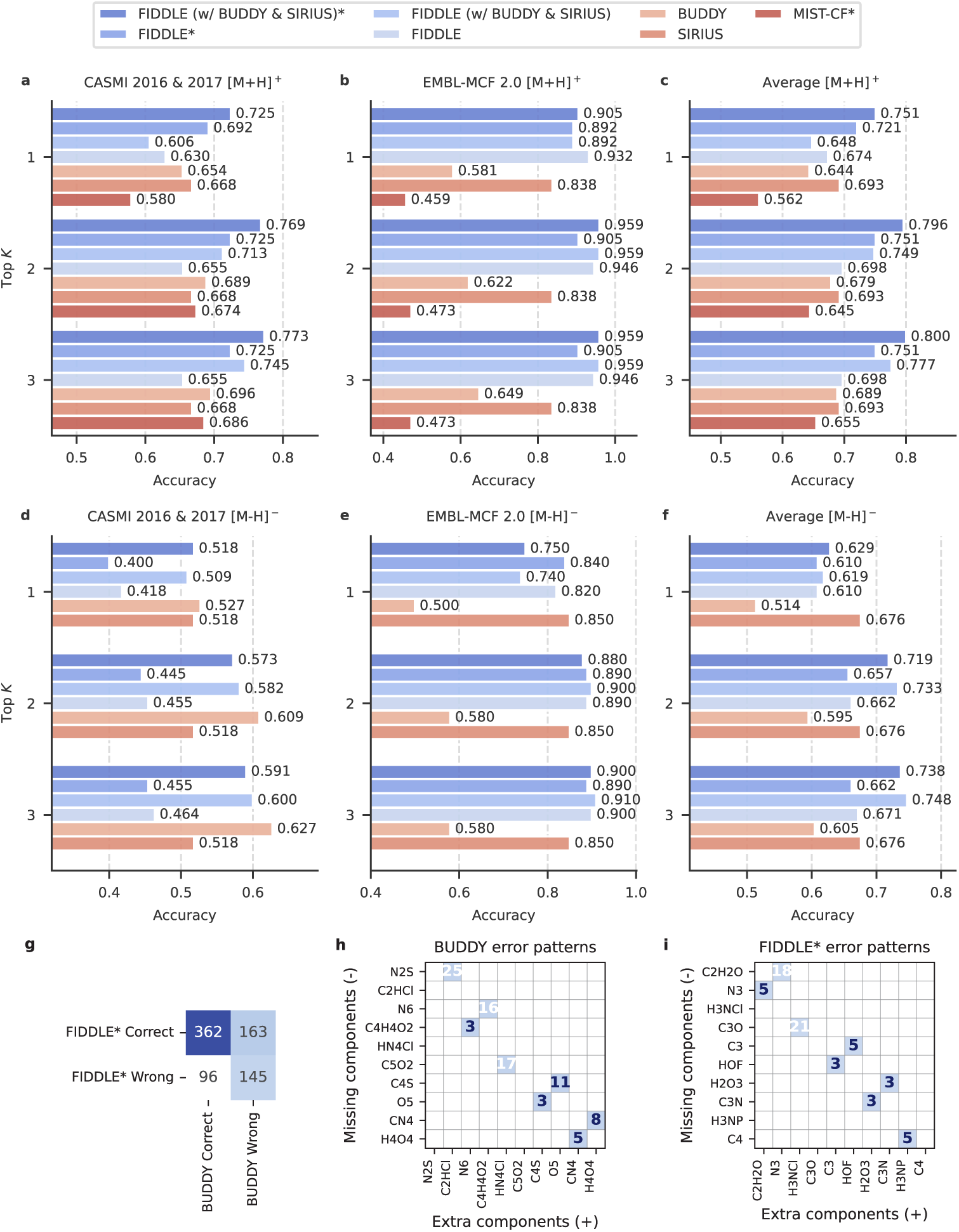
Performance and error patterns of formula identification on external test sets. Methods marked with an asterisk (*) denote deep learning models trained on all available datasets, whereas models without an asterisk were trained on all available datasets except for the NIST23 dataset. Following the original configuration of MIST-CF, we only evaluated the data acquired under the positive ion mode.

## 3 Discussion

In this work, we introduce a deep learning approach named FIDDLE to identify chemical formulas from tandem mass spectra. It consists of three steps: predicting formulas from tandem mass spectra, generating candidate formulas, and ranking the candidates based on predicted confidence scores. FIDDLE not only accelerates the formula identification compared to state-of-the-art algorithms (BUDDY and SIRUIS), but also outperforms them on both the evaluation set (with 88.3% top-1 accuracy and 93.6% top-5 accuracy) and external benchmarking data sets (with the average top-3 accuracy of 80.0% and 73.8% for positive and negative ion modes, respectively, across three datasets). Our noise robustness evaluation on 1,000 test spectra shows that FIDDLE maintains over 90% accuracy even at large noise levels, significantly outperforming BUDDY and SIRIUS on both Q-TOF and Orbitrap data. A separate evaluation on chimeric spectra confirmed that FIDDLE’s performance decreases as spectral mixing increases; this finding, while expected, will guide future design of data augmentation strategies to enhance its training process.

According to our ablation study, both data augmentation and contrastive loss between manually combined spectra pairs enhance formula identification from tandem mass spectra. The post-processing steps, which include generating candidate formulas and ranking them by predicted confidence scores, aim to refine the predicted formulas using the SENIOR rules. These steps refine most candidate formulas with no more than three missed heavy atoms, and significantly alleviate the challenge of incorrect hydrogen counts due to their small mass. Furthermore, because FIDDLE and other formula identification methods perform well on different compounds and spectra, combining their predicted candidate formulas and ranking them by using FIDDLE’s confidence score prediction further improves the accuracy of the predicted formulas, even though its running time may be longer. This suggests that a promising direction for future work is the development of hybrid methods that unify the predictive power of deep learning with the systematic rigor of search-based approaches.

We note that, even though the contrastive loss is employed to take into account the effect of experimental conditions, this effect is not completely eliminated, as FIDDLE still shows variations in accuracy under different conditions. Future work may be focused on improving accuracy across different conditions, especially those with less training data. The workflow of FIDDLE offers several avenues, such as improving MS/MS representations by pre-trained models from self-supervise learning [30], using MS/MS prediction methods for data augmentation [31], and extending the workflow with adduct type prediction for comprehensive application.

Furthermore, instead of simply accepting all top K candidates, confidence score thresholds could be used to implement false discovery rate (FDR) control. This approach would introduce a critical standard from proteomics to the field of metabolomics, where such controls remain largely unexplored.

Beyond formula identification, FIDDLE serves as a robust foundation for compound structural elucidation from MS/MS spectra, enabling the inference of covalent bonds between atoms based on their atomic composition. In the future, this methodology could be expanded to characterize diverse molecular structures by leveraging machine learning techniques to extract structural information from MS/MS spectra and integrating this information into molecular structure characterization.

## 4 Methods

### 4.1 MS/MS data pre-processing

#### 4.1.1 MS/MS data filtering

The training and testing MS/MS datasets are compiled from several sources, including NIST20, NIST23, MoNA, GNPS, Agilent PCDL, and an internal dataset, following the preprocessing methodology described by Hong et al. [31]. To construct the internal dataset, we acquired 1,424 compounds from APExBIO [32] and measured their tandem mass spectra using a Waters Synapt G2 mass spectrometer at 40 eV with various precursor types, including [M + H]^+^, [M + Na]^+^, [2 M + H]^+^, [M + 2 H]^2+^, and [M – H]^-^. To minimize the interference from impurities in the air, e.g., water vapor (H_2_O) and carbon dioxide (CO_2_), we excluded peaks below 50 *m/z* from the scans. Additionally, three benchmarking sets generated for community-wide evaluation of metabolite identification algorithms are used, including CASMI 2016, CASMI 2017, and EMBL-MCF 2.0.

These datasets undergo a series of filtering steps to ensure data quality: 1) Mass spectra with fewer than five peaks are excluded due to potential unreliability. 2) The *m/z* range is confined to (0, 1500] to account for the rarity of spectra with *m/z* values above 1500. 3) Only the molecules composed of frequent atoms (C, H, O, N, F, S, Cl, P, B, I, Br, Na, and K) are retained. (iv) Only spectra associated with common precursor types are included, e.g. [M + H]^+^, [M + Na]^+^, [2 M + H]^+^, [M + H – H_2_O]^+^, [M + H – 2 H_2_O]^+^, [M + H – NH_3_]^+^, [M + H + NH_3_]^+^, [M + H – CH_2_O_2_]^+^, [M + H – CH_4_O_2_]^+^ for positive modes, and [M – H]^-^, [2 M – H]^-^, [M – H – CO_2_]^-^, [M – H – H_2_O]^-^ for negative modes, along with [M + 2 H]^2+^, a doubly charged precursor type. 4) The total count of atoms in the molecules is capped at 300 to exclude the molecules not typically classified as ‘small molecules’. 5) The tolerance for precursor mass discrepancy is set at 10 ppm for Q-TOF and 5 ppm for Orbitrap instruments, ensuring precise mass matching. The statistics information of the datasets are detailed in the following Table 1. The specific instruments contained in each instrument type are shown in Extended Table 1.

#### 4.1.2 Precursor *m/z* simulation

The precursor *m/z* from NIST20, NIST23, CASMI 2016, and CASMI 2017 are theoretical values so they are adjusted via random shifts, following the approach used by Xing et al. [16] and the observations by Böcker at el. [17] that the mass deviations fit a Gaussian distribution with the standard deviation of 1*/*3 of the mass tolerance. We sampled the deviations from Gaussian distributions within the set tolerance ranges (5 ppm for Orbitrap and 10 ppm for Q-TOF) to simulate the experimental conditions accurately. These simulated precursor *m/z* values are utilized throughout both the training and testing phases, enhancing the model’s generalizability for application in real-world formula identification tasks.

#### 4.1.3 Simplification of precursor types

As depicted in Fig. 2 (g), the dataset exhibits significant imbalance across different precursor types. To improve formula identification for less common precursor types, we simplify the precursor types by eliminating uncharged molecules, such as water (H_2_O), ammonia (NH_3_), carbon dioxide (CO_2_), formic acid (CH_2_O_2_), and acetic acid (CH_4_O_2_), adding them into the original formula representation (for the detailed precursor types see Extended Table 2). For example, consider a molecular formula C_6_H_12_N_2_O_2_ with the precursor type [M + H + NH_3_]^+^. After simplifying the precursor type, the formula is adjusted to C_6_H_15_N_3_O_2_, and the precursor type is simplified to [M + H]^+^, reflecting the integration of NH_3_ into the molecular formula. This simplification is recorded allowing the predicted formulas to be converted back to the original formula with the original precursor type. Through this process, many uncommon precursor types are consolidated into more common precursor types, which expand the training data of common precursor types, thereby enhancing the predictions.

#### 4.1.4 Training set construction and data augmentation

Each dataset, either acquired using Q-TOF or Orbitrap MS/MS instruments, is first split into training and test sets according to their molecular canonical SMILES, which ensures that there are no common compounds between the training set and the test set. It is worth noting that canonical SMILES cannot distinguish stereoisomers, so this splitting strategy guarantees that stereoisomers with different configurations are not separated between training and test sets. Two strategies were employed for data splitting: 1) random splitting (see the model in Section 2.4 and the model marked with an asterisk in Section 2.5); 2) leaving spectra of unique compounds from NIST23 for evaluation (see the model in Section 2.2 and the model without an asterisk in Section 2.5). Then, for contrastive learning (CL), we constructed MS/MS spectra pairs from the training sets. The spectra are grouped by canonical SMILES. For each compound, we randomly picked another compound from the same group to construct a positive pair and another compound from a different group to construct a negative pair.

To enhance the robustness of the deep learning model, we generated augmented MS/MS spectra by perturbing the spectra in the training set. Specifically, we added random noise sampled from a Gaussian distribution with a mean of 0 and a standard deviation of 0.1 (∼ N (0, 0.1)) to the intensities of an experimental Q-TOF spectrum, generating two augmented spectra for each Q-TOF spectrum and thus tripling the size of the training set from Q-TOF mass spectrometry. On average, the cosine similarity between the augmented spectrum and the corresponding experimental spectrum is 0.936, which is close to the similarity between of replicated spectra of the same compound (0.977) in the Q-TOF training set. However, noise adding lead to significantly lower similarity for Orbitrap spectra (as shown in Extended Fig. 4). Since the training size for Orbitrap is sufficiently large, we did not apply data augmentation for training deep learning model for Orbitrap.

### 4.2 Representation learning for MS/MS spectra

#### 4.2.1 Model architecture

The representation learning for MS/MS spectra is structured as a two-stage process: MS/MS embedding and the elimination of the experimental condition effect. This approach decomposes the complex task of MS/MS representation learning under multiple experimental conditions into more tractable steps, facilitating the network’s ability to extract meaningful features. Additionally, the resultant condition-independent MS/MS features reduce the complexity of the decoder.

Acknowledging the significance of the correlation among fragment ions in MS/MS, such as the neutral loss between two ions, dilated convolutions with large kernels are employed, building upon the previous work for *de novo* peptide sequencing [22]. Due to the large effective receptive field (ERF), this method allows the deep neural network to learn meaningful patterns across fragment ions with long mass ranges from the MS/MS data. Each convolutional block consists of two sequentialized dilated convolutions with ReLU activation, random dropout, a residual connection, and maxpooling. The max-pooling uses a kernel size of 2 and a stride size of 2, which halves the spectra length. Upsampling along the channel dimension is applied in the skip connection when the output and input channel numbers differ. Dilated convolutions with exponentially increasing dilation factors (1, 2, 4, 8, etc.) and large kernels rapidly expand the ERF without significantly increasing the number of parameters [21, 33]. We can calculate the ERF for our model as:

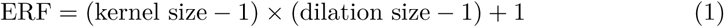

where kernel sizes are 45, 43, 41, 39, 37, and 35, and the dilation sizes are 1, 2, 4, 8, 8, and 8, respectively. The total ERF of this model is 1153, indicating that each position in the MS/MS feature is directly influenced by 1153 input bins. It is worth noting that while the ERF defines the range of direct input influence, the model can still capture patterns beyond this range. In addition, we utilize weight normalization to speed up the convergence and reduce the dependency of normalization on batch size, enabling training with small batches and limited GPU memory [34].

The features from all the blocks are concatenated together along the channel dimension, resulting in a 1024 × 235 tensor, where 1024 results from the stacked convolutional blocks with channel sizes of 32, 32, 64, 128, 256, and 512, and 235 is the length of the MS/MS spectra after pooling layers. Then, through a global pooling layer along the spectra-length dimension, the tensor is pooled into a 1024-dimensional vector as the embedded MS/MS feature. Precursor types, normalized collision energy, and simulated precursor *m/z* are included as experimental conditions. These conditions are embedded into a 16-dimensional vector through linear layers. The embedded MS/MS feature and experimental conditions are concatenated and fed into sequential linear layers for the condition-independent MS/MS features. These features are constrained by contrastive learning so that features from the same molecule are learned to be the same, while features from different molecules are learned to be different.

#### 4.2.2 Loss function

Given a pair of binned spectra *i* and *j*, the encoder of the prediction model embeds them into condition-independent MS/MS feature vectors **z_i_** and **z_j_**. These feature vectors are constrained by the contrastive loss:

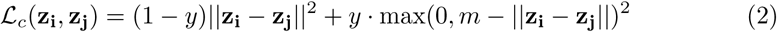

where *y* is the label between the paired spectra. If spectra *i* and *j* are from the same molecule, the label is 1; otherwise, the label is 0. *m* is a hyperparameter (by default, *m* = 1.0) that defines the lower bound distance between spectra of different molecules. Three auxiliary tasks, including the atom count prediction, the molecular mass prediction, and the prediction of the hydrogen-to-carbon (H/C) ratio, are used for model regularization to improve the model generalizability [24]. Mean squared error (MSE) losses are applied to these tasks. The loss function for sample *i* from the pair (*i, j*) is defined as:

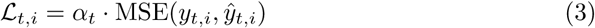

where ℒ*_t,i_* denotes the loss for task *t* on sample *i*; *y_t,i_* and *ŷ_t,i_* denote the label and prediction, respectively; and *α_t_* denotes the weight of loss on task *t*. In the experiments, we use the weights of 0.01, 1, and 10 for these three auxiliary tasks, respectively. Finally, the loss function for training the model is:

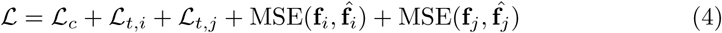

### 4.3 Formula refinement algorithm

The candidate formula refinement algorithm (Algorithm 1) refines an input chemical formula into a list of formulas that comply with the SENIOR rules [25] and match a target precursor mass within a specified tolerance. This algorithm can start with the formula predicted by the deep learning model or a formula identified by tools such as SIRIUS or BUDDY, facilitating the integration of results from multiple methods. The algorithm initializes lists to track refined formulas, previously explored formulas, and current candidates with their associated search depths. By iteratively modifying the counts of heavy atoms without exceeding a set search depth, the algorithm systematically refines each candidate formula. During the exploration, one heavy atom is added or removed, and hydrogen counts are adjusted to align with the precursor mass. The refinement process concludes either when a predefined number of formulas have been explored or when all candidate formulas are exhausted, incorporating a timeout check. The algorithm ultimately produces a list of formulas that align closely with the target precursor mass and adhere to the SENIOR rules.

#### Algorithm 1 Candidate Formula Refinement Algorithm

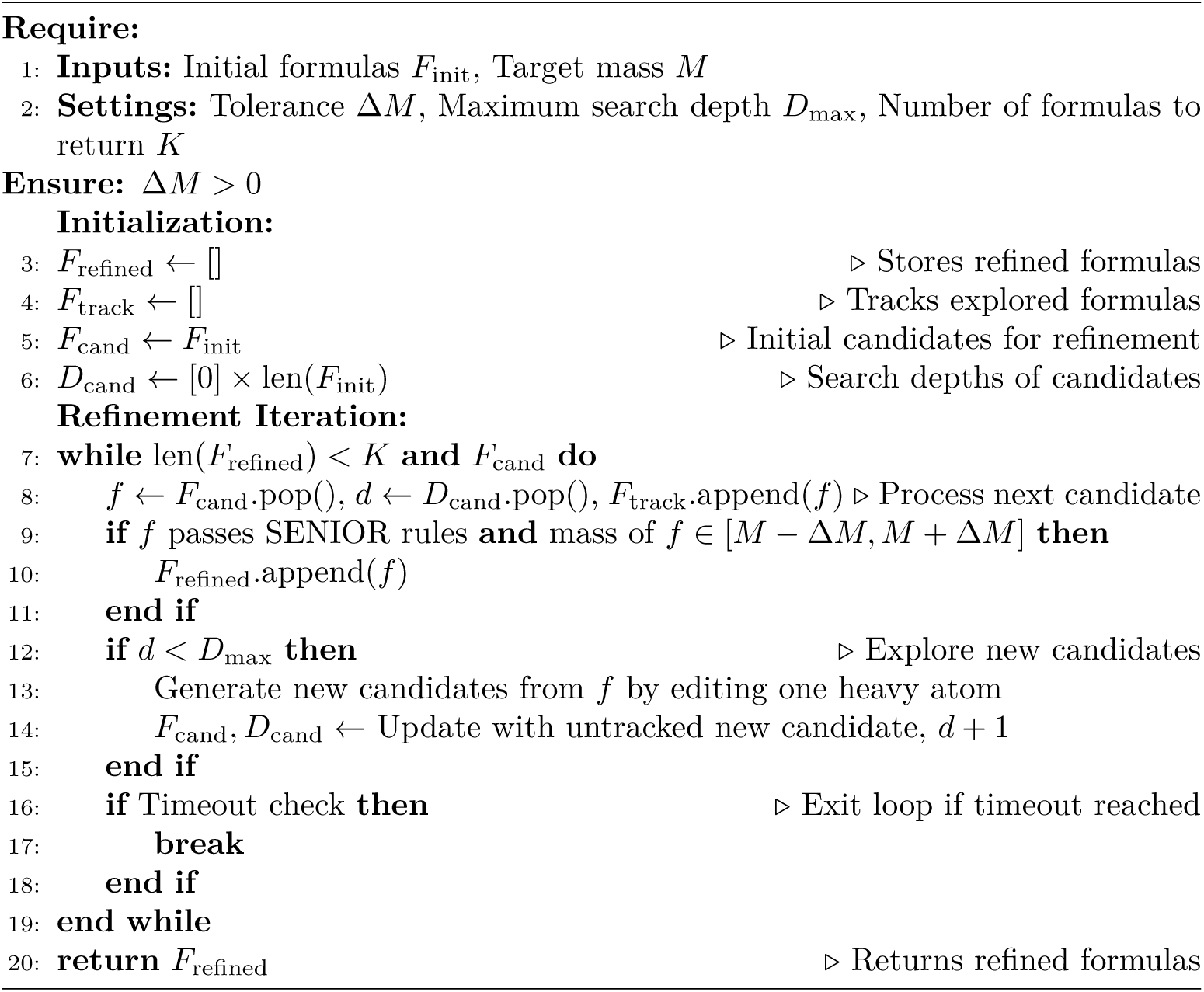

### 4.4 Prediction of confidence score

The inputs for the confidence score prediction model consist of the concatenated condition-independent MS/MS feature vector and the vector representation of the predicted formula. The model architecture comprises a sequence of five linear layers, each followed by batch normalization, ReLU activation function, and random dropout for regularization. The dimensions of the five layers are 416, 208, 104, 26, and 13, respectively. A Sigmoid activation function is used in the final layer to produce a confidence score between 0 and 1 with higher values indicating more likely correct formula assignments. Training data for this model is obtained from the inference results of initial formula identification followed by candidate formula refinement. Specifically, each of the top 5 refined candidate formulas is labeled, with 0 indicating a correct candidate and 1 denoting an incorrect one. Since this model has significantly fewer trainable parameters than the formula prediction model, the training set is randomly sampled down to 10,000 instances if it exceeds this limit to prevent overfitting.

### 4.5 Implementation and computing environment

Both deep neural networks (formula prediction and candidate scoring) were implemented in PyTorch version 1.13.1, comprising 21,027,933 (about 21M) and 20,055,938 (about 20.1M) parameters, respectively. A list of specific parameter numbers for each layer/block is described in Supplementary Table 1 and 2.

The training of models in FIDDLE was conducted using a batch size of 512 over 200 epochs, with an initial learning rate of 0.001. The models were optimized using AdamW and a ReduceLROnPlateau learning rate schedule that reduces the learning rate when a metric has stopped improving for 5 epochs.

The training device was two NVIDIA RTX A6000 GPUs equipped with 48 GB of memory. For inference only, the minimum GPU memory requirement is 500 MB when using a batch size of 1. The deep learning framework used was PyTorch version 1.13.0, running on Ubuntu 20.04.1.

To facilitate broader adoption within the research community, we release our implementation as an open-source Python package (msfiddle) with optimized routines for both GPU and CPU-only environments, enabling reproducible access to pre-trained models and inference pipelines with standardized APIs compatible with common scientific computing workflows.

### 4.6 Running SIRIUS and BUDDY

We installed SIRIUS v6.0.1, last updated on July 21st, 2024, for the Linux (64-bit) platform, accessible at https://github.com/boecker-lab/sirius. Additionally, the BUDDY v0.3.6 PyPI package, updated on January 29th, 2024, was obtained from https://github.com/Philipbear/msbuddy. All experiments were performed on a workstation with 20.04.1-Ubuntu 64-bit operating system equipped with 16 Intel(R) Core(TM) i7-9800X CPUs. For comprehensive configuration details, please refer to Extended Table 3).

## 5 Data availability

NIST20 and NIST23 libraries can be accessed with a license from NIST23 - Mass Spectral Library. Data from GNPS can be downloaded from GNPS - Mass Spectral Libraries. The MoNA, CASMI16, CAMI17, and EMBL-MCF 2.0 libraries can be downloaded from Mass Bank of North America. The Agilent PCDL is available from Agilent Technologies (https://www.agilent.com). The Waters Q-TOF dataset can be obtained by contacting us.

## 6 Code availability

The source code for all experiments is publicly available at GitHub: https://github.com/JosieHong/FIDDLE under the CC BY-NC-SA 4.0 License. Pre-trained models for Q-TOF and Orbitrap MS/MS data can be downloaded from GitHub release: https://github.com/JosieHong/FIDDLE/releases. A command line tool msfiddle is available via PyPI (https://pypi.org/project/msfiddle), providing simple usage to download, manage, and use models.

## Acknowledgements

We acknowledge the Center for Bioanalytical Metrology (CBM), an NSF Industry-University Cooperative Research Center, for providing funding under grant NSF IIP-1916645. This work was also partially supported by National Science Foundation grant DBI-2011271.

## 7 Extended Data

**Extended Table 1:**
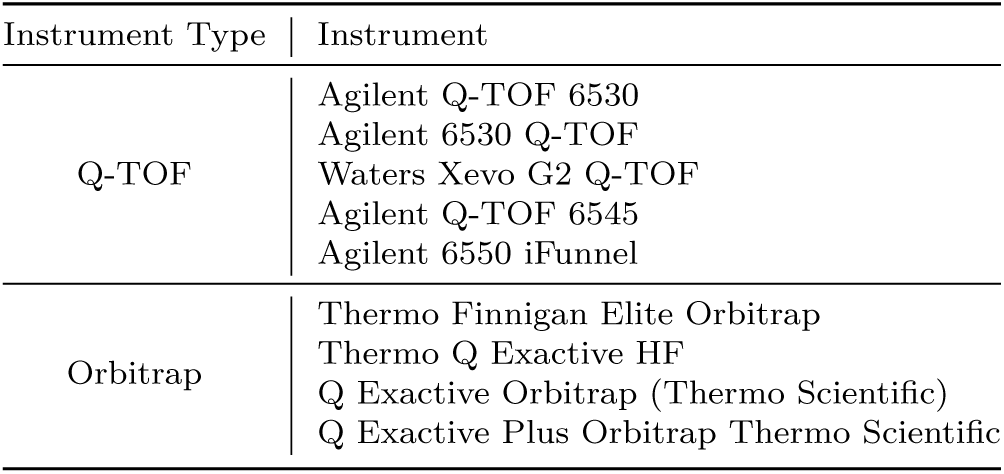
List of Q-TOF and Orbitrap instruments.

| Instrument Type | Instrument |
| --- | --- |
| Q-TOF | Agilent Q-TOF 6530 |
|  | Agilent 6530 Q-TOF |
|  | Waters Xevo G2 Q-TOF |
|  | Agilent Q-TOF 6545 |
|  | Agilent 6550 iFunnel |
| Orbitrap | Thermo Finnigan Elite Orbitrap |
|  | Thermo Q Exactive HF |
|  | Q Exactive Orbitrap (Thermo Scientific) |
|  | Q Exactive Plus Orbitrap Thermo Scientific |

**Extended Table 2:**
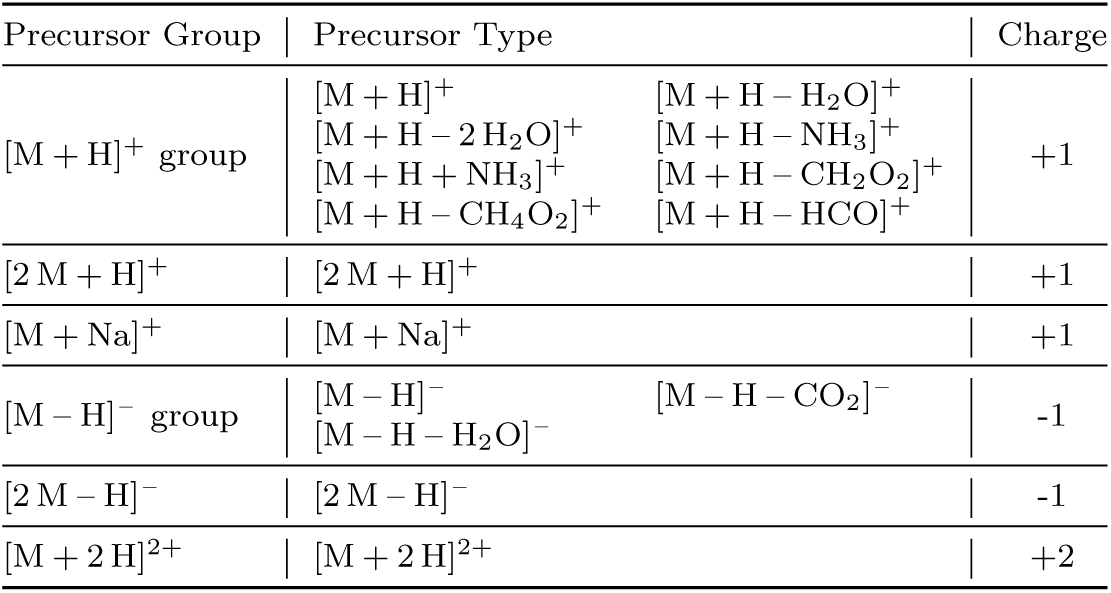
List of simplified precursor types.

**Extended Table 3:** Configurations of SIRIUS and BUDDY.

**Extended Table 2:** List of simplified precursor types.
|  |  |  |  |
| --- | --- | --- | --- |
| 1151 |  |  |  |
| 1152 |  | SIRIUS | BUDDY |
| 1153 | Version | 6.0.1 | 0.3.6 |
| 1154 |  |  |  |
| 1155 | Atom Type Limitation | H, C, N, O, P, B,<br>Cl, Br, S, F, I | H, C, N, O, P, B,<br>Cl, Br, S, F, I |
| 1156 |  |  |  |
| 1157 | | | $[M + H]^+$, $[M + H - NH_3]^+$, |
| 1158 | | | $[M + H + NH_3]^+$, $[M + H + H_2O]^+$, |
| 1159 | Precursor Type Limitation | $[M + H]^+$, $[M + H - H_2O]^+$,<br>$[M + H - CH_4O_2]^+$,<br>$[M + H - H_4O_2]^+$,<br>$[M - H]^-$, $[M - H - H_2O]^-$ | $[M + H - H_2O]^+$, $[M + H - H_2O]^+$,<br>$[M + H - HCO]^+$,<br>$[M + K]^+$, $[M + Na]^+$,<br>$[M - H]^-$, $[M - H + CH_2O_2]^-$,<br>$[M - H + CH_4O_2]^-$,<br>$[M + Cl]^-$, $[M + Br]^-$ |
| 1160 |  |  |  |
| 1161 |  |  |  |
| 1162 |  |  |  |
| 1163 | Charge Limitation | +1, -1 | +1, -1 |
| 1164 | Timeout for Each Compound | 300 s | 600 s |
| 1165 |  |  |  |
| 1166 | Mass Deviation Tolerance | 10 ppm | 10 ppm |

**Extended Table 4:** Configurations for training MIST-CF.

**Extended Table 3:** Configurations of SIRIUS and BUDDY.
|  |  |  |
| --- | --- | --- |
| 1171 |  |  |
| 1172 | **Data Preprocessing** |  |
| 1173 | Candidate Selection | “COMMON” filter [15] and FastFilter [19] |
| 1174 |  |  |
| 1175 | Atom Type Limitation | H, C, N, O, P, B,<br>Cl, Br, S, F, I,<br>Si, Se, Fe, Co, As, Na, K |
| 1176 |  |  |
| 1177 | Precursor Type Limitation | $[M + H]^+$, $[M + H - H_2O]^+$, $[M + H + NH_3]^+$,<br>$[M + H - H_4O_2]^+$, $[M + Na]^+$, $[M + K]^+$ |
| 1178 |  |  |
| 1179 | Charge Limitation | +1 |
| 1180 | Mass Deviation Tolerance | 10 ppm |
| 1181 | Maximum Number of Peaks for Subformula Assignment | 20 |
| 1182 | Intensity Threshold of Peaks (Normalized to 1) | 0.003 |
| 1183 |  |  |
| 1184 | **Model Architecture** |  |
| 1185 | Number of Layers | 2 |
| 1186 | Dropout Rate | 0.1 |
| 1187 | Hidden Layer Dimensionality | 128 |
| 1188 | Fourier Feature Representation | Absolute Sine Features |
| 1189 | **Training Parameters** |  |
| 1190 | Batch Size | 4 |
| 1191 | Maximum Number of Decoys | 32 |
| 1192 | Maximum Number of Epochs | 200 |
| 1193 | Learning Rate | 0.00045 |
| 1194 | Learning Rate Decay Factor | 0.88 |

**Extended Fig. 1:**
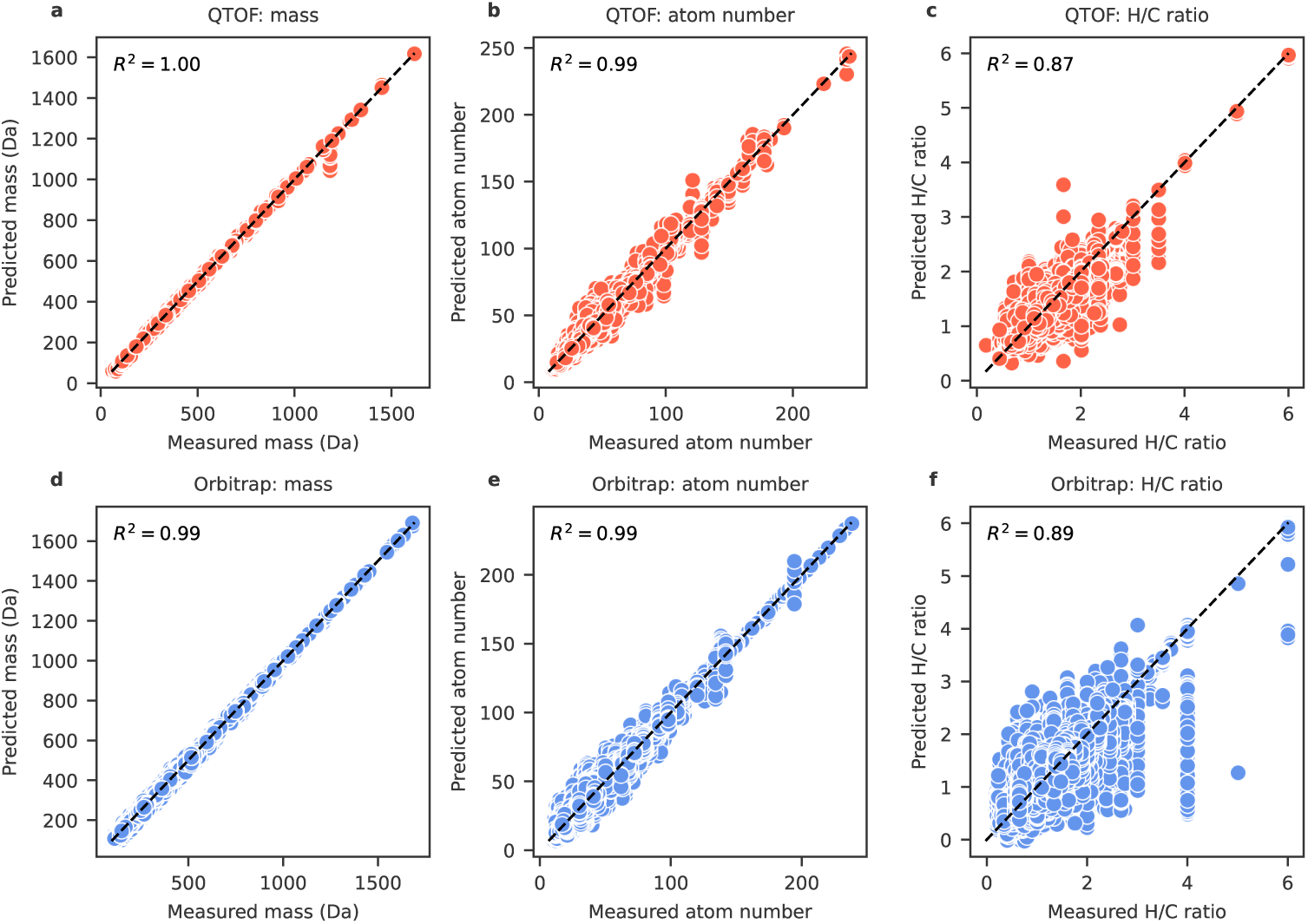
Performance on auxiliary tasks, including mass, atom number, and H/C ratio prediction. (a), (b), and (c) correspond to Q-TOF MS/MS, while (d), (e), and (f) correspond to Orbitrap MS/MS.

**Extended Fig. 2:**
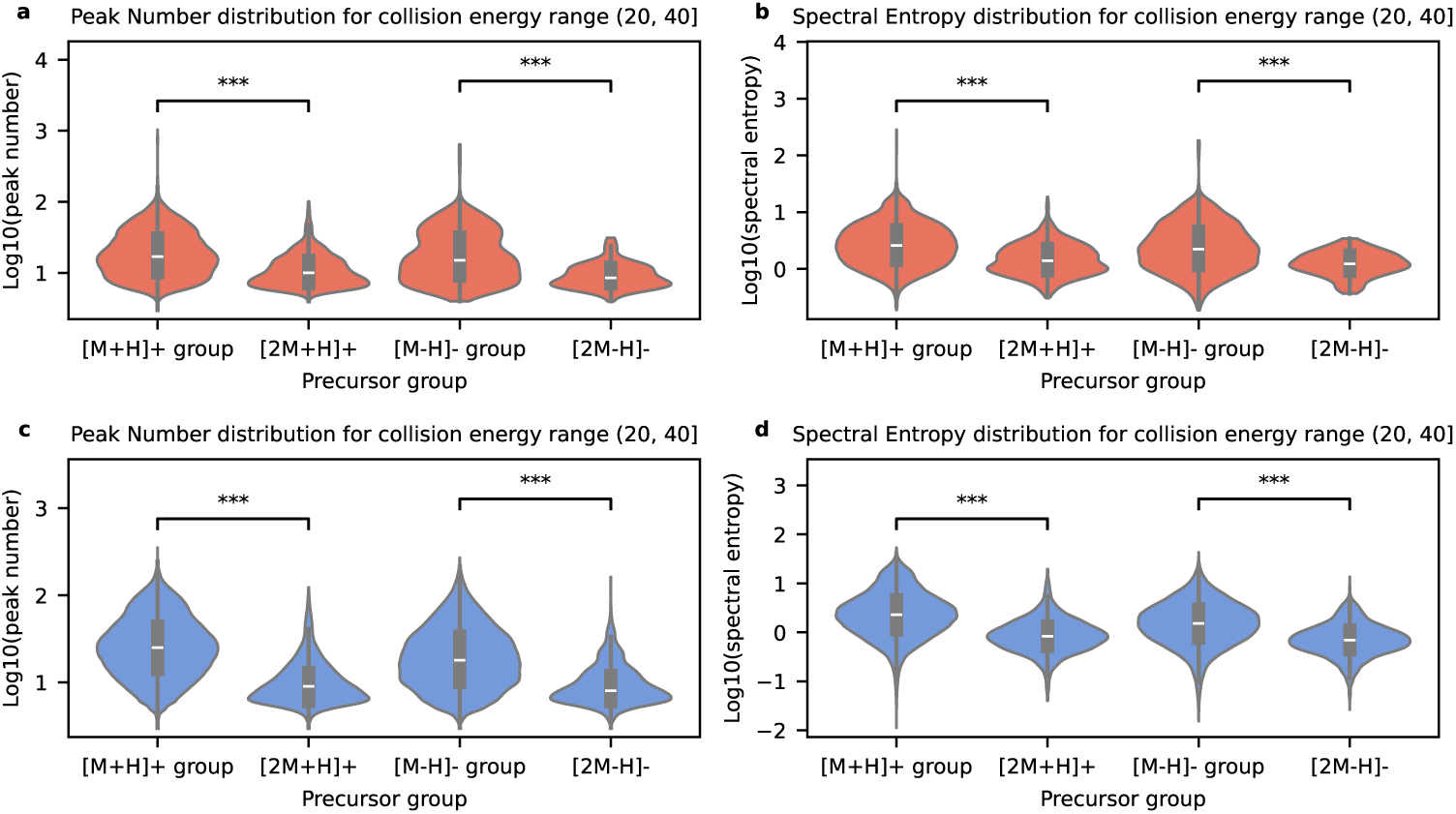
Number of peaks and spectral entropy distributions. The spectra under collision energy range (20, 40] serve as an example. Spectra in other collision energy ranges reveal that [M + H]^+^ and [M – H]^-^ groups contain more complex spectra compared to [2 M + H]^+^ and [2 M – H]^-^ groups. The asterisk (***) denotes in the Mann-Whitney U test the p-value is smaller than 0.001, so the two hues are statistically significantly different.

**Extended Fig. 3:**
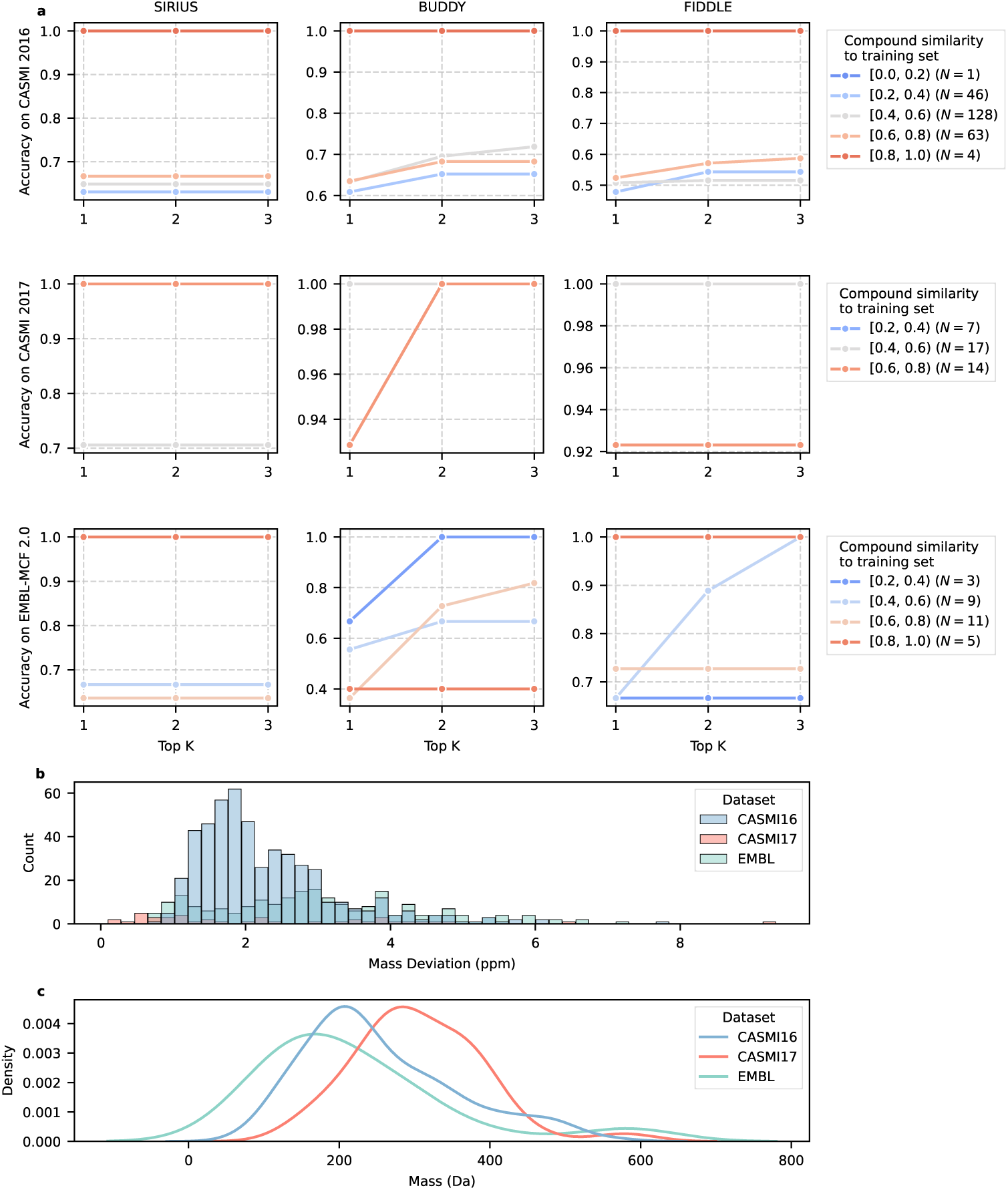
Potential impacts on performance on external test sets. (a) Performance on compounds with varying similarities to the training set. For each compound in the test set, the maximum Tanimoto similarity with any compound in the training set is used as its compound similarity to the training set. FIDDLE’s performance is correlated with training set compound similarity, indicating its purely data-driven approach learns patterns without incorporating prior knowledge. (b) and (c) show the distributions of mass deviation and molecular mass in the test sets, respectively. It is worth noting that although the CASMI datasets were collected using Q-TOF instruments, we simulated the precursor *m/z* values as described in Section 4.1.2, resulting in a lower mass deviation compared to the EMBL dataset.

**Extended Fig. 4:**
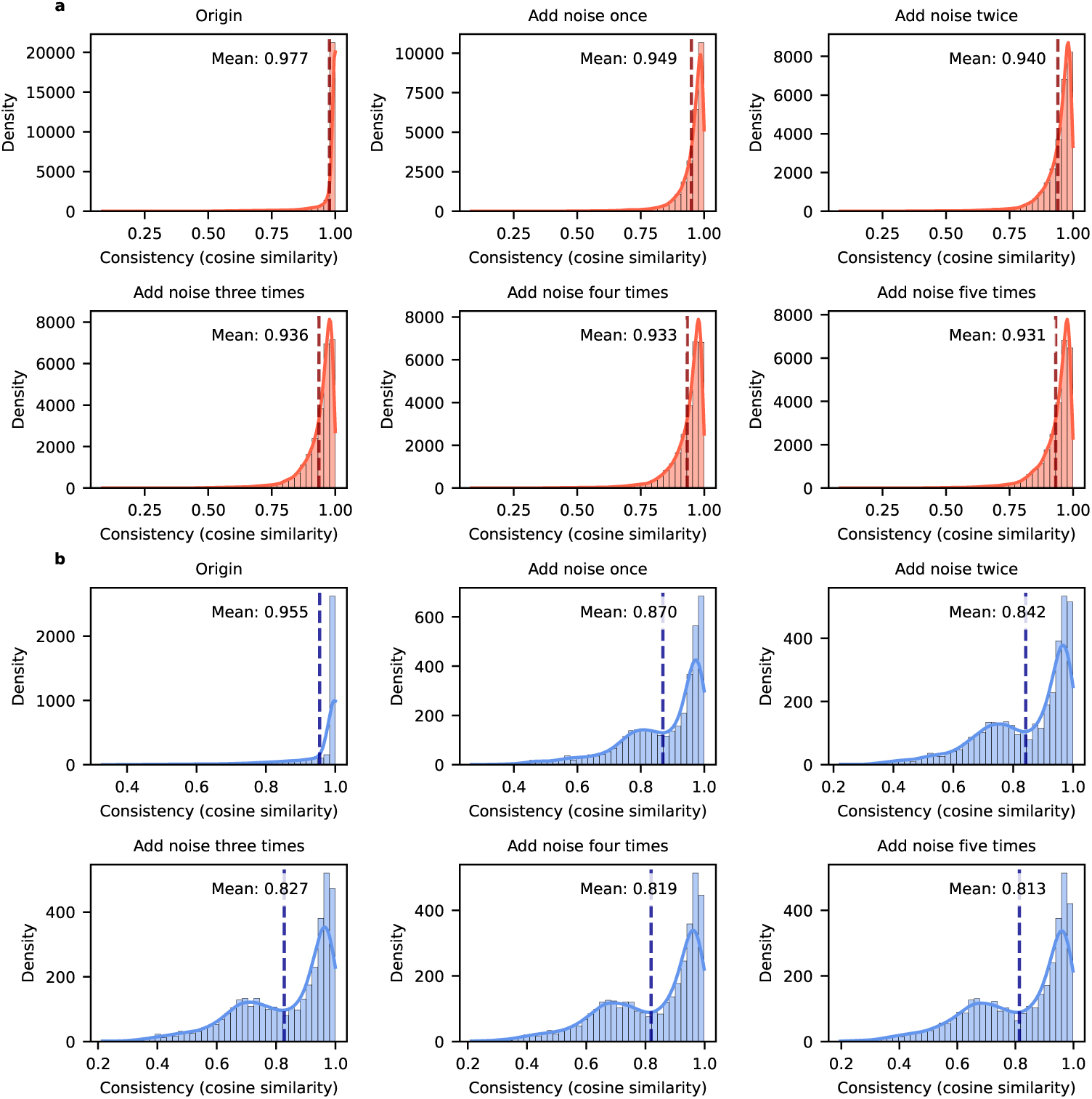
Consistency before and after data augmentation. (a) and (b) correspond to Q-TOF and Orbitrap, respectively. Note that data augmentation was not applied to Orbitrap in the experiments, as adding noise significantly decreases the MS/MS consistency for Orbitrap, and the training size for Orbitrap is quite large.

## Notes

https://github.com/JosieHong/FIDDLE

https://pypi.org/project/msfiddle/

### Competing Interest Statement

The authors have declared no competing interest.

### Summary of Updates

Ablation studies added to quantify contributions of data augmentation, contrastive loss, and post-processing (Fig. 4e f g). Robustness evaluation included with varying noise levels and chimeric spectra, showing superior stability (lines 311, 325). Data splitting analysis expanded with formula-based partitions to prevent structural similarity between train and test (Supplementary Fig. 3). Case studies added comparing predictions from BUDDY and FIDDLE, highlighting complementary error patterns (Fig. 5g h i). Representation space analysis performed, linking latent diversity with prediction accuracy (Fig. 3e). Compound class performance evaluated across 17 chemical superclasses, identifying strengths and weaknesses (Fig. 3f).

