## Supplementary Information for "FIDDLE: a deep learning method for chemical formulas prediction from tandem mass spectra"

### 1 Supplementary figures and tables

#### 1.1 Supplementary Fig. 1 | percentage of double-charged spectra

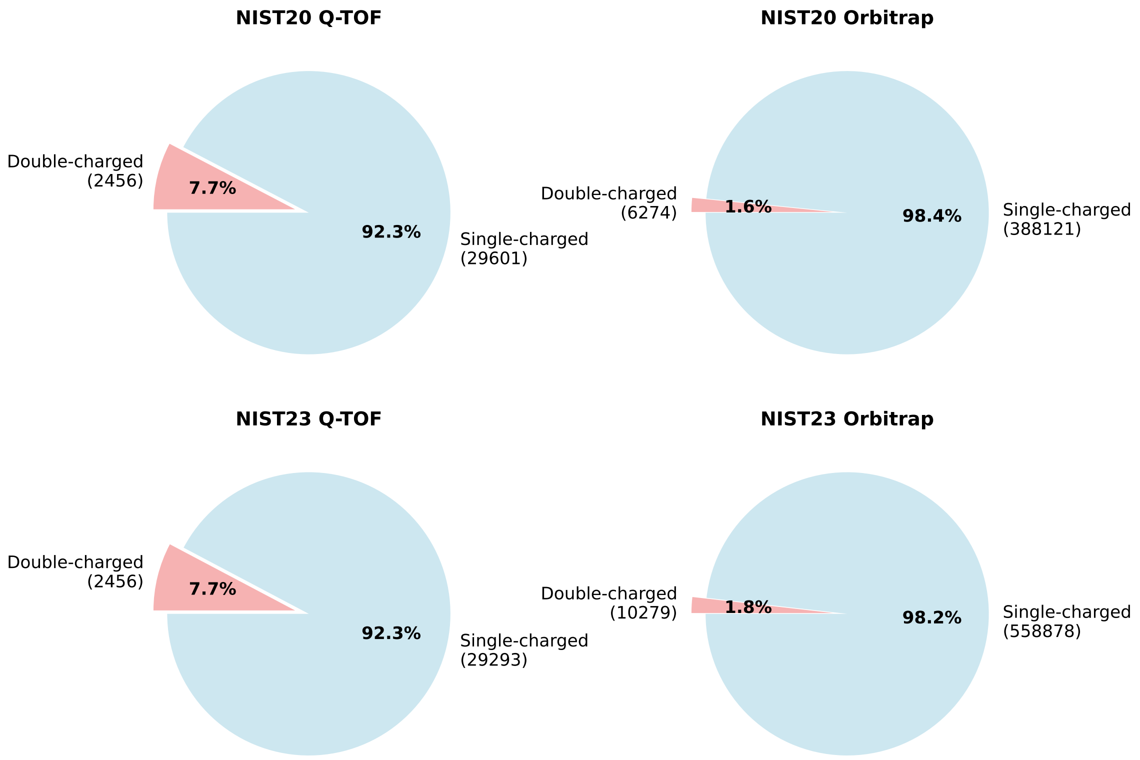

Supplementary Fig. 1: **Percentage of double-charged spectra in NIST20 and NIST23 datasets**, which are not supported by SIRIUS [3]. Other datasets contain no double-charged spectra after preprocessing.

#### 1.2 Supplementary Fig. 2 | BUDDY formula coverage analysis

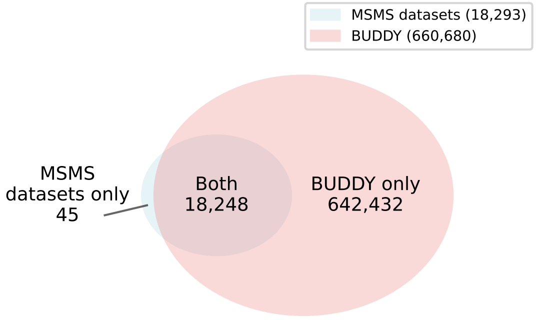

(a)

| | |
|---|---|
| $C_{42}H_{71}N_5O_{29}$ | $C_{72}H_{101}N_{17}O_{26}$ |
| $C_{12}H_{17}N_4OS$ | $B_2C_{20}H_{21}NO_5$ |
| $BC_7H_7O_4$ | $C_{74}H_{120}O_{38}$ |
| $C_{73}H_{131}N_3O_{31}$ | $C_{29}F_7H_{29}N_4O_9$ |
| $BC_{12}H_{11}O_3$ | $BC_3H_5N_2O_2$ |
| $BC_{14}H_{16}NO$ | $C_{44}H_{72}N_6O_{32}$ |
| $BC_8H_9O_4$ | $C_{14}H_{17}N_2O_4$ |
| $BC_5H_6NO_2$ | $C_{76}H_{104}N_{18}O_{19}S_2$ |
| $C_{48}H_{81}N_5O_{33}$ | $C_{75}H_{118}N_{20}O_{22}S$ |
| $C_7H_9N_2O$ | $C_{71}H_{112}O_{37}$ |
| $C_{29}H_{48}N_6O_{19}$ | $BC_{12}FH_{10}O_2$ |
| $C_{76}H_{118}N_{16}O_{19}$ | $C_{38}H_{64}N_6O_{26}$ |
| $C_{54}H_{96}N_4O_{17}S$ | $C_{69}H_{108}N_5O_{34}P$ |
| $C_{71}H_{110}N_{24}O_{18}S$ | $BC_5FH_5NO_2$ |
| $BC_5H_5O_4$ | $BBrC_6H_6O_2$ |
| $BC_7H_7O_3$ | $BC_4H_7N_2O_2$ |
| $C_{26}H_{45}N_3O_{20}$ | $C_{74}H_{120}O_{39}$ |
| $BC_7H_{10}NO_4$ | $BC_7H_9O_3$ |
| $C_{34}H_{58}N_4O_{24}$ | $BC_5H_5O_3S$ |
| $BC_6ClH_6O_2$ | $C_{20}H_{19}N_5Na_2O_6$ |
| $C_{47}H_{76}N_6O_{35}$ | $C_{73}H_{118}O_{38}$ |
| $C_{44}H_{74}N_6O_{31}$ | $BC_6H_8NO_2$ |
| $BC_5H_{12}NO_4S$ | |

(b)

Supplementary Fig. 2: **BUDDY formula coverage analysis** (a) and 45 uncovered molecular formulas (b). The coverage is analyzed by searching all unique formulas from preprocessed datasets using `mass_to_formula` implemented in `msbuddy` 0.3.12 with 10 ppm tolerance, where halogens are included.

#### 1.3 Supplementary Fig. 3 | comparison of splitting strategies

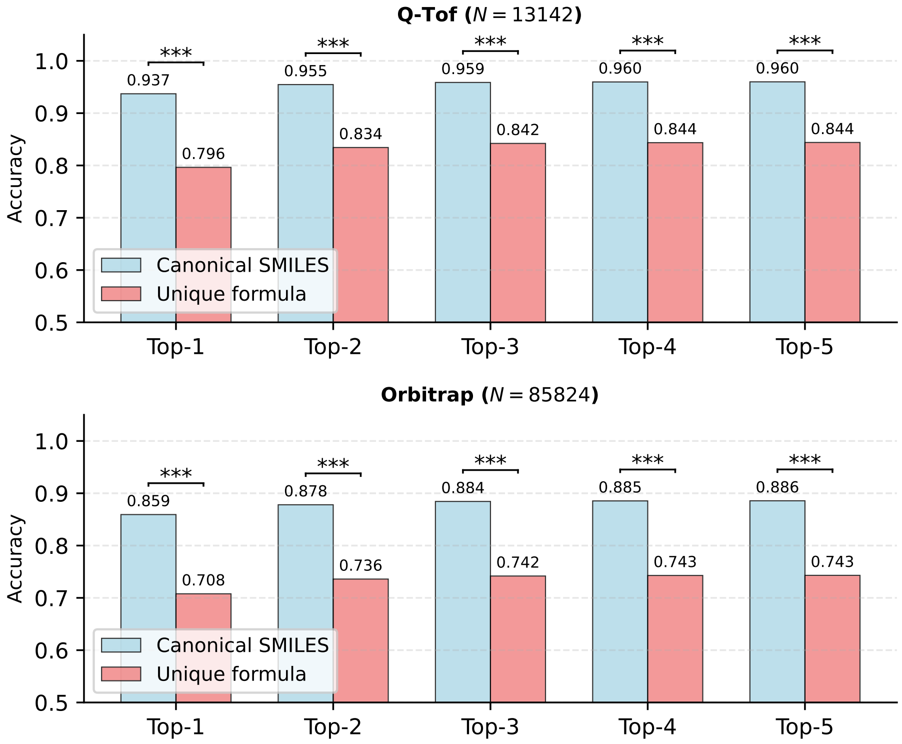

Supplementary Fig. 3: **Comparison of splitting strategies with FIDDLE**. Blue and red bars represent accuracy on overlapping samples from splits based on canonical SMILES and unique molecular formula, respectively. One-sample t-tests indicate all $p$-values are below 0.001, annotated as ***.

#### 1.4 Supplementary Fig. 4 & 5 | top-5 accuracy on external test sets

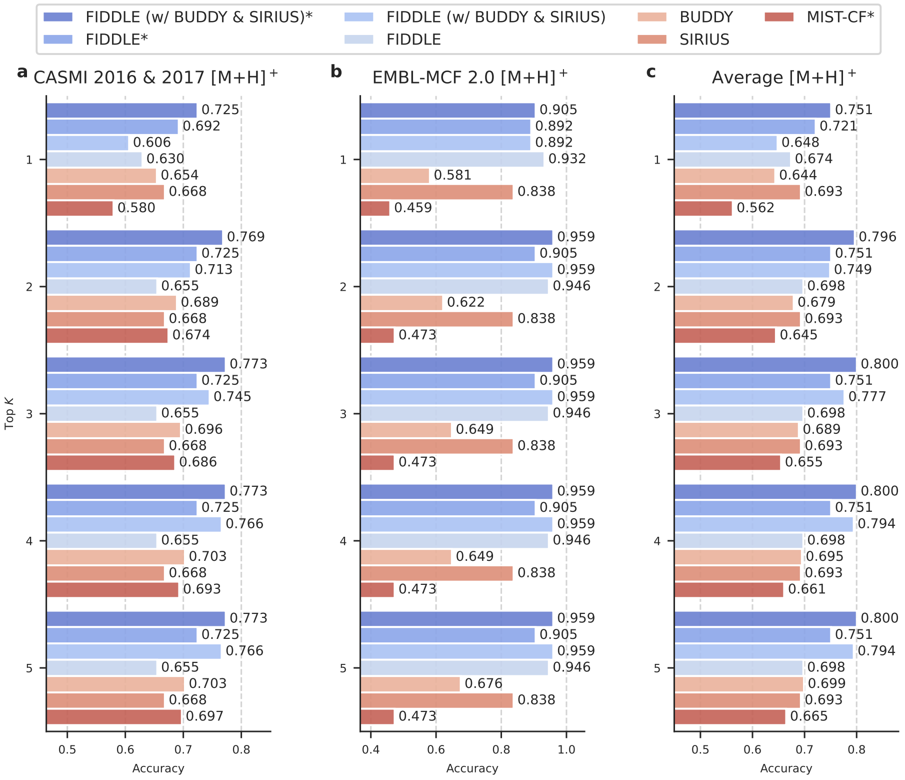

Supplementary Fig. 4: Top-5 accuracy of formula identification on external test sets (positive ion mode). Methods marked with an asterisk (*) denote deep learning models trained on all available datasets, while unmarked models exclude the NIST23 dataset. MIST-CF was evaluated only in positive ion mode following its original configuration.

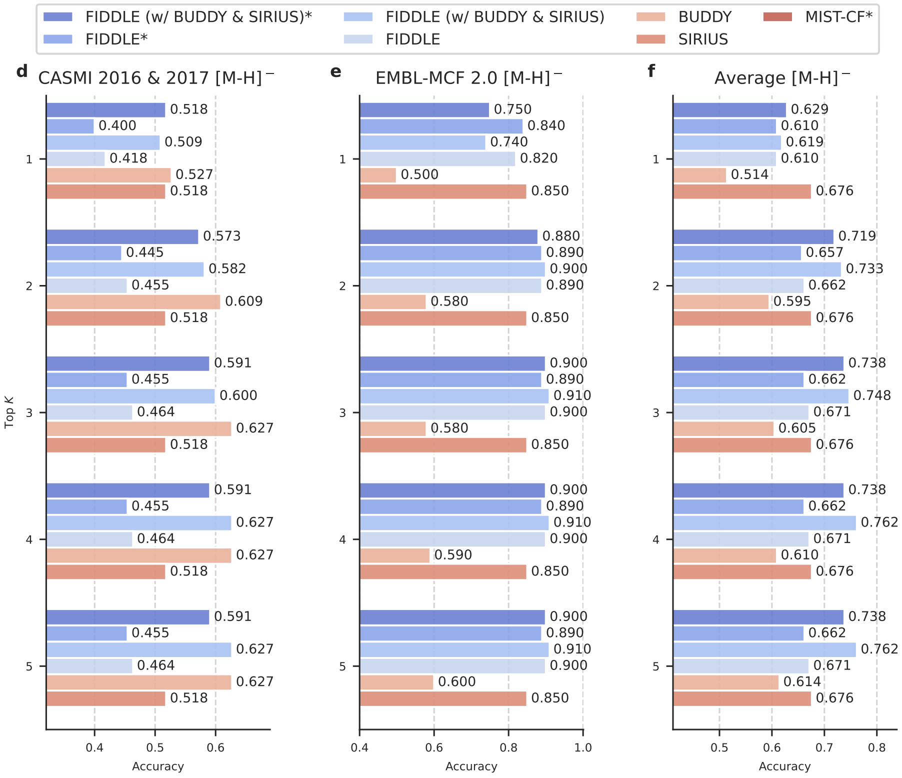

Supplementary Fig. 5: Top-5 accuracy of formula identification on external test sets (negative ion mode). Methods marked with an asterisk (*) denote deep learning models trained on all available datasets, while unmarked models exclude the NIST23 dataset. MIST-CF was evaluated only in positive ion mode following its original configuration.

#### 1.5 Supplementary Table 1 & 2 | parameter breakdown for FIDDLE

| **Component** | **Parameters** | **Percentage** |
|---|---|---|
| *TCN Encoder* | | |
| Block 1 (1→32, k=45) | 47,744 | 0.2% |
| Block 2 (32→32, k=43) | 88,192 | 0.4% |
| Block 3 (32→64, k=41) | 254,336 | 1.2% |
| Block 4 (64→128, k=39) | 967,424 | 4.6% |
| Block 5 (128→256, k=37) | 3,671,552 | 17.5% |
| Block 6 (256→512, k=35) | 13,896,704 | 66.1% |
| *TCN Subtotal* | *18,925,952* | *90.1%* |
| *Metadata Embeddings* | | |
| Mass embedding (1→4) | 12 | 0.0% |
| Collision energy embedding (1→4) | 12 | 0.0% |
| Adduct type embedding (7×4) | 28 | 0.0% |
| *Embedding Subtotal* | *52* | *0.0%* |
| *Feature Fusion* | | |
| FC1 (1036→512) | 531,456 | 2.5% |
| FC2 (512→512) | 263,168 | 1.3% |
| *Feature Fusion Subtotal* | *794,624* | *3.8%* |
| *Decoders* | | |
| Formula decoder | 330,070 | 1.6% |
| Mass decoder | 325,745 | 1.5% |
| Atom number decoder | 325,745 | 1.5% |
| H/C ratio decoder | 325,745 | 1.5% |
| *Decoder Subtotal* | *1,307,305* | *6.2%* |
| **Total Parameters** | **21,027,933 (21.0M)** | **100.0%** |

Supplementary Table 1: **Parameter numbers and percentage of the formula prediction model in FIDDLE.**

| **Component** | **Parameters** | **Percentage** |
|---|---:|---:|
| *TCN Encoder* | | |
| Block 1 (1→32, k=45) | 47,744 | 0.2% |
| Block 2 (32→32, k=43) | 88,192 | 0.4% |
| Block 3 (32→64, k=41) | 254,336 | 1.3% |
| Block 4 (64→128, k=39) | 967,424 | 4.8% |
| Block 5 (128→256, k=37) | 3,671,552 | 18.3% |
| Block 6 (256→512, k=35) | 13,896,704 | 69.3% |
| *TCN Subtotal* | *18,925,952* | *94.4%* |
| *Metadata Embeddings* | | |
| Mass embedding (1→4) | 12 | 0.0% |
| Collision energy embedding (1→4) | 12 | 0.0% |
| Adduct type embedding (7×4) | 28 | 0.0% |
| *Embedding Subtotal* | *52* | *0.0%* |
| *Feature Fusion* | | |
| FC1 (1036→512) | 531,456 | 2.6% |
| FC2 (512→512) | 263,168 | 1.3% |
| *Feature Fusion Subtotal* | *794,624* | *4.0%* |
| *FDR Decoder* | | |
| Layer 1 (525→416) | 219,232 | 1.1% |
| Layer 2 (416→208) | 86,944 | 0.4% |
| Layer 3 (208→104) | 21,840 | 0.1% |
| Layer 4 (104→52) | 5,512 | 0.0% |
| Layer 5 (52→26) | 1,404 | 0.0% |
| Layer 6 (26→13) | 364 | 0.0% |
| Final layer (13→1) | 14 | 0.0% |
| *FDR Decoder Subtotal* | *335,310* | *1.7%* |
| **Total Parameters** | **20,055,938 (20.1M)** | **100.0%** |

Supplementary Table 2: **Parameter numbers and percentage of the candidate scoring model in FIDDLE.**

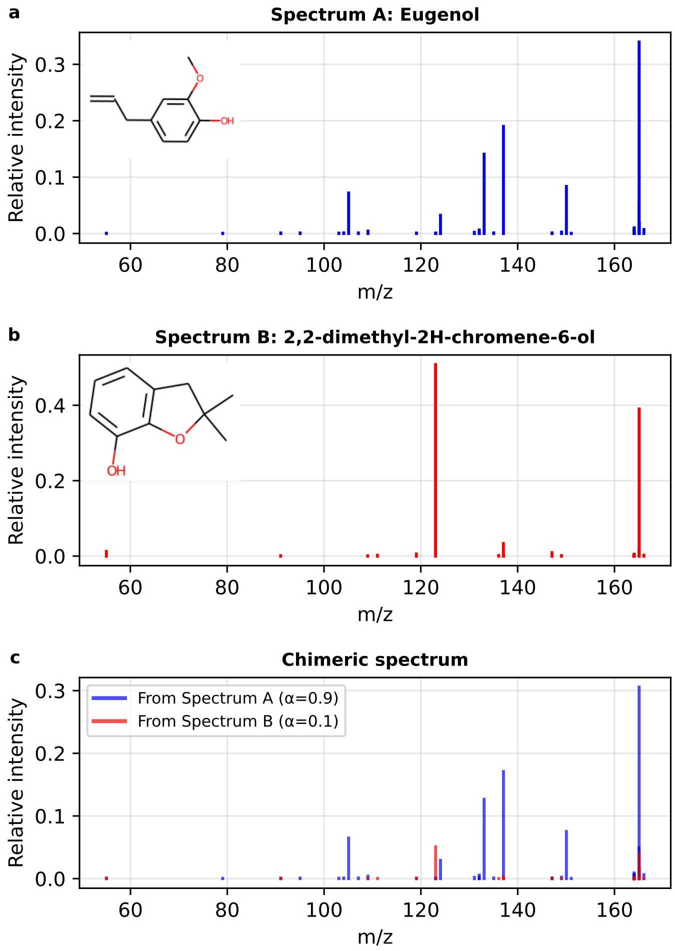

Supplementary Fig. 6: **An example of synthetic chimeric spectra**, where the weight ($\alpha$) is 0.9 for spectrum A, and 0.1 for spectrum B.

### 2 Supplementary notes

#### 2.1 Supplementary Note 1 | evaluation on chimeric spectra

Real-world MS/MS spectra from complex samples are often chimeric, containing fragments from multiple co-isolated precursors that hinder accurate formula identification [5]. Because our existing datasets consist exclusively of single-compound spectra, we constructed a synthetic chimeric dataset to evaluate performance under these more challenging conditions. To create this dataset, we performed a weighted merge of spectra from pairs of different compounds that were both mass-similar (mass difference ($< 0.005 Da$) and acquired under identical experimental conditions (collision energy and precursor type).

Figure Supplementary Fig. 6 illustrates an example chimeric spectrum created by merging Eugenol and 2,2-dimethyl-2H-chromene-6-ol with mixing ratios of $\alpha = 0.9$ and $\alpha = 0.1$, respectively,

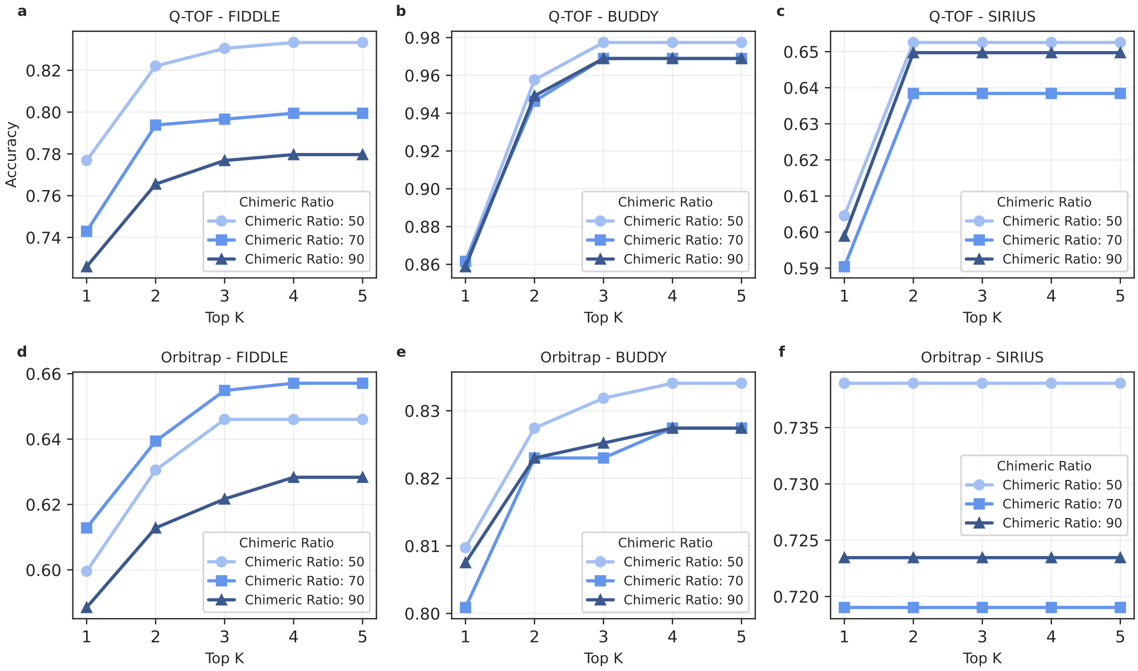

Supplementary Fig. 7: **Performance on synthetic chimeric spectra.**

where spectral features from both compounds contribute to the synthetic chimeric spectrum. In these experiments, chimeric spectra were constructed for three mixing ratios ($\alpha = 0.5, 0.7, 0.9$) on both Q-TOF and Orbitrap data, yielding 354 and 452 spectra for each choice of mixing ratio, respectively.

For evaluation, we used the formula of the compound with the larger $\alpha$ value as the ground truth label for each chimeric spectrum; however, for spectra with $\alpha = 0.5$ (equal mixing), formulas from either compound were considered correct predictions.

The results (Supplementary Fig. 7) show that FIDDLE's performance degrades significantly on chimeric spectra for both Q-TOF and Orbitrap data, as it was not designed for or trained on such data. In contrast, BUDDY and SIRIUS demonstrate greater robustness because their methods do not rely on the fine-grained MS/MS features learned by deep neural networks. Interestingly, most methods achieved optimal performance at an equal mixture ratio ($\alpha = 0.5$), where either precursor could be considered the ground truth. The sole exception was FIDDLE on Orbitrap data. Future enhancements to FIDDLE could address this limitation by incorporating synthetic chimeric spectra during training or by developing more generalizable deep learning architectures.

#### 2.2 Supplementary Note 2 | compounds classification

The test compounds are classified by ClassyFire [2], which automatically classifies chemical compounds into a comprehensive taxonomy of over 4,800 categories across 11 hierarchical levels (Kingdom, SuperClass, Class, SubClass, etc.) based solely on their molecular structures. In the experiments, we use ClassyFire Batch, which retrieves and formats ClassyFire results for InChIKey identifiers.

| Index | Superclass |
|---|---|
| 0 | Organophosphorus compounds |
| 1 | Organic 1,3-dipolar compounds |
| 2 | Alkaloids and derivatives |
| 3 | Lipids and lipid-like molecules |
| 4 | Lignans, neolignans and related compounds |
| 5 | Phenylpropanoids and polyketides |
| 6 | Nucleosides, nucleotides, and analogues |
| 7 | Organic acids and derivatives |
| 8 | Organic oxygen compounds |
| 9 | Organic nitrogen compounds |
| 10 | Hydrocarbon derivatives |
| 11 | Organoheterocyclic compounds |
| 12 | Benzenoids |
| 13 | Organosulfur compounds |
| 14 | Organic Polymers |
| 15 | Organohalogen compounds |
| 16 | Unknown |

Supplementary Table 3: Index and Superclass from ClassyFire

Notably, we identified a bug in the ClassyFire Batch implementation where the system stops processing InChIKeys mid-query when the status shows "Completed" but no results are returned for kingdom, superclass, or other taxonomic levels[1]. To address this issue, we manually restarted the classification process multiple times whenever the processing terminated.

In the end, all the compounds are classified into 16 superclasses as shown in Supplementary Table 3, where the compounds with no results are denoted as 'Unknown'. We chose the superclass as the taxonomic level for performance analysis because it provides an optimal balance: higher levels (such as kingdom) are too broad to meaningfully distinguish between different types of compounds, while lower levels (such as subclass or molecular framework) create categories that are too granular, resulting in classes with insufficient sample sizes for statistically meaningful analysis.

#### 2.3 Supplementary Note 3 | analysis of spectral representation space

We analyzed the MS/MS spectral representation space in FIDDLE by applying the t-SNE dimensionality reduction to project the 512-dimensional latent vectors into a 2D visualization, as shown in Supplementary Fig. 8 (Q-TOF) and Supplementary Fig. 9 (Orbitrap). The heatmap colors indicate the prediction accuracy by FIDDLE on the test set in Supplementary Fig. 8 (a), and the diversity of formulas in the training set in (c), respectively, across different regions in the representation space. Here, the formula diversity is computed as the average Euclidean distance between pairs of formula vectors within each cell of the heatmap. For cells with over 1000 formula pairs, a random sample of 1000 pairs is used for computing efficiency. The observed patterns confirm a correlation between MS/MS representations and formula diversity, which shows that regions with similar spectral representations but highly divergent chemical formulas pose significant challenges for accurate formula prediction by FIDDLE, because FIDDLE did not learn distinctive features in the representation space that can distinguish these chemical formulas. This formula diversity typically

[1] https://bitbucket.org/fiehnlab/classyfire-batch/issues/11/classyfire-batch-stops-processing

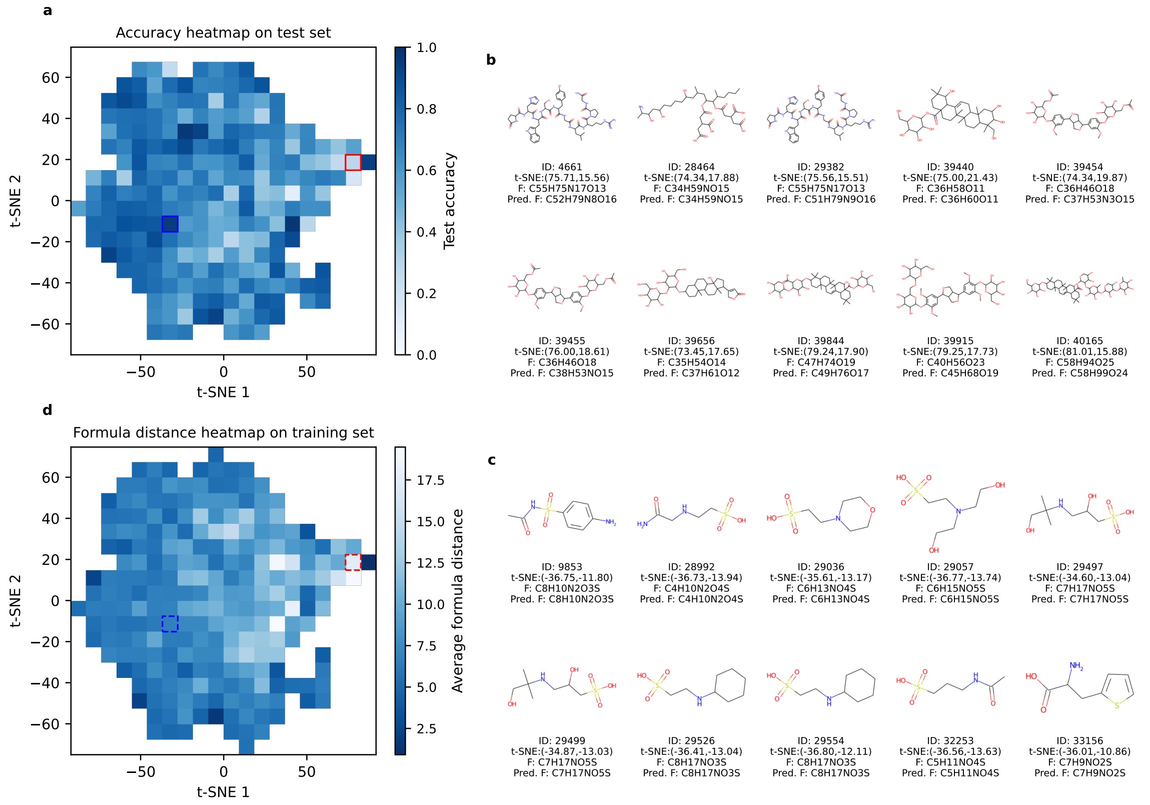

Supplementary Fig. 8: **Visual analysis of Q-TOF MS/MS spectral representation space**. Heatmaps generated using t-SNE illustrate FIDDLE's prediction accuracy on the test set (a) and the diversity of formulas in the training set (c). Some compounds with poor prediction accuracy (within the red box in (a)) are shown in (b), in comparison with some well-predicted compounds (within the blue box in (a)).

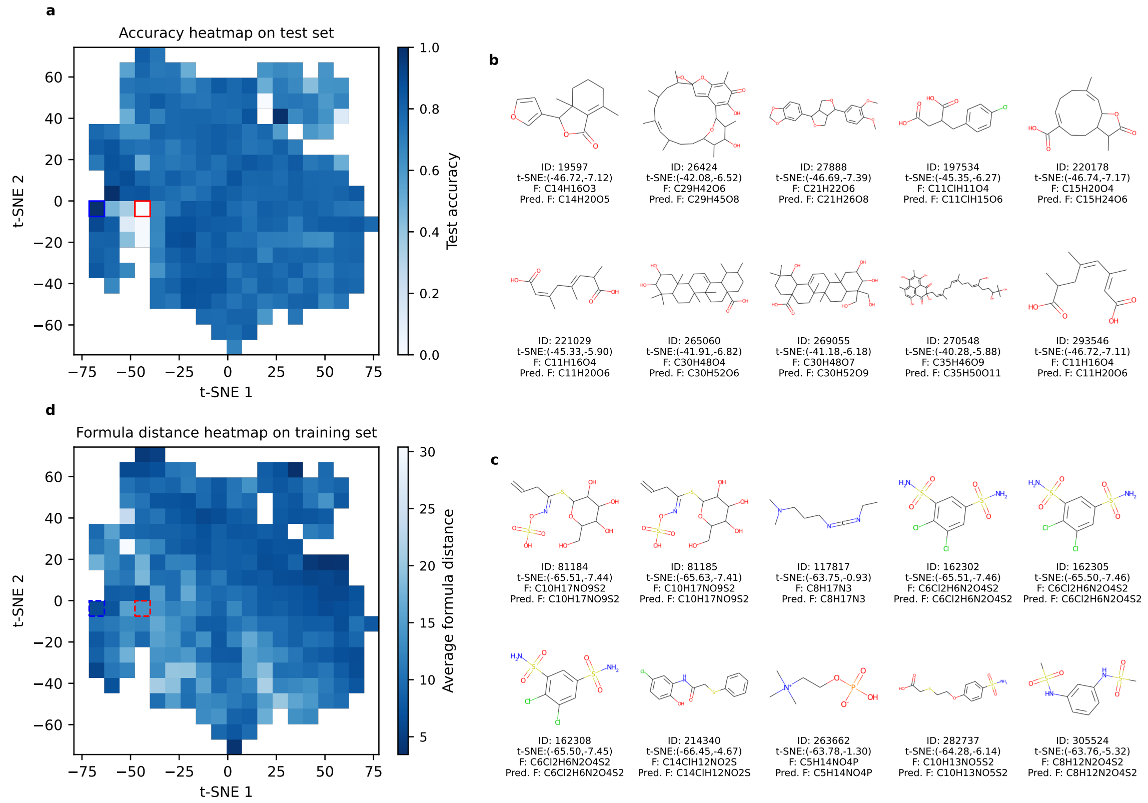

Supplementary Fig. 9: **Visual analysis of Orbitrap MS/MS spectral representation space**. Heatmaps generated using t-SNE illustrate FIDDLE's prediction accuracy on the test set (a) and the diversity of formulas in the training set (c). Some compounds with poor prediction accuracy (within the red box in (a)) are shown in (b), in comparison with some well-predicted compounds (within the blue box in (a)).

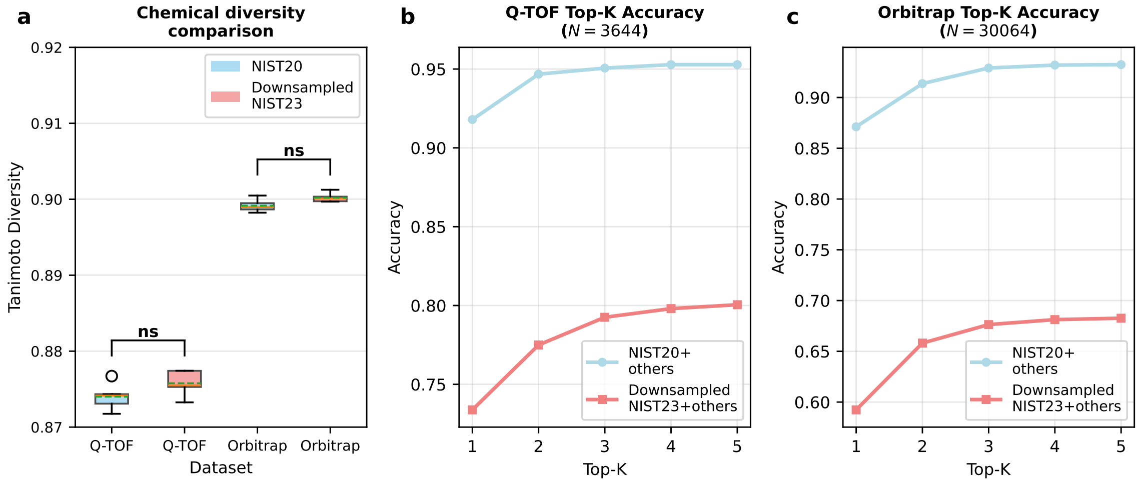

Supplementary Fig. 10: **Performance comparison of FIDDLE trained on NIST20 versus downsampled NIST23**, both combined with additional datasets and evaluated on the same randomly split test set from the additional datasets. The number of spectra in the test sets for Q-TOF and Orbitrap are $N = 32,281$ and $N = 269,113$, respectively.

arises from increased atom counts rather than more complex atomic compositions, as evidenced by the fact that 100% grid cells contain molecules with identical atomic compositions for Q-TOF and Orbitrap data. Similar patterns are observed in the Orbitrap results.

Supplementary Fig. 8 (b) illustrates that structurally distinct molecules can produce remarkably similar MS/MS patterns, despite having different molecular formulas ($F$). For example, spectra 39454 ($C_{36}H_{46}O_{18}$) and 39656 ($C_{35}H_{54}O_{14}$) exhibit proximal representations in the t-SNE space, even though their formulas are quite different. This phenomenon can occur when diverse molecular scaffolds yield overlapping fragment ion patterns, often driven by structural features like glycosidic bonds, ester linkages, and heteroatom-rich functional groups. These features, while informative, can complicate formula prediction. Future improvements to FIDDLE could benefit from a deeper investigation of these challenging cases. In contrast, Supplementary Fig. 8 (d) demonstrates that regions of low formula diversity, where fragmentation patterns are more closely tied to specific structural motifs, facilitate accurate formula prediction. In these instances, unique spectral features enable clear differentiation between molecular formulas.

#### 2.4 Supplementary Note 4 | analysis of contributions by NIST23

We systematically evaluated whether NIST23's contribution to performance improvement stems from the dataset size by comparing size-matched training scenarios: NIST20 with other datasets (Agilent PCDL, MoNA, GNPS, and Waters Q-TOF) versus downsampled NIST23 (matched to NIST20's size) combined with the identical other datasets. Chemical diversity was quantified through average pairwise Tanimoto similarities [1] of 2,000 randomly sampled compounds (5 independent iterations). Supplementary Fig. 10 reveals that while both configurations show statistically equivalent chemical diversity (panel a), reducing NIST23 to match NIST20's size results in significant performance degradation across all top-K accuracy metrics ($K = 1, 2, ..., 5$) (panels b-c). This clear relationship between dataset size reduction and performance loss demonstrates that NIST23's advantages are

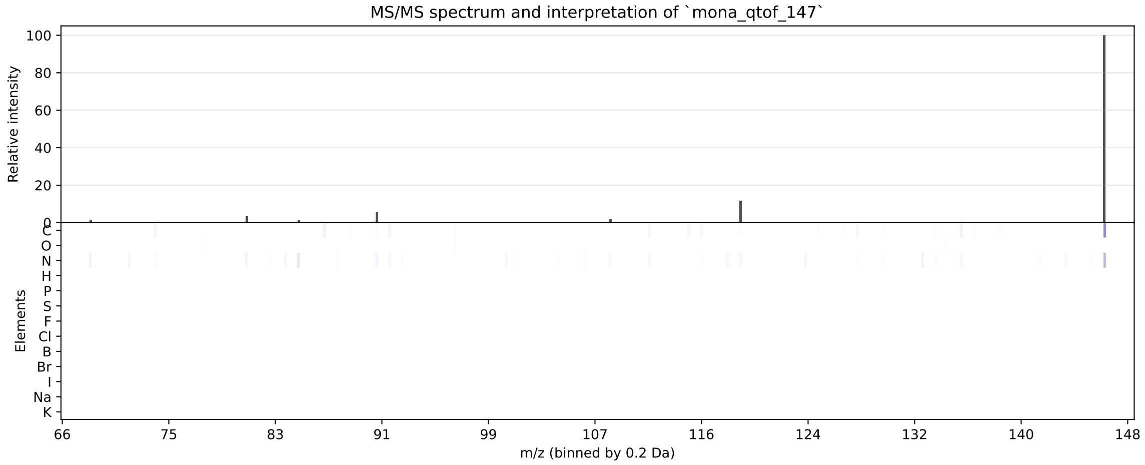

Supplementary Fig. 11: **MS/MS spectra and Shapley values showing peak contributions to atomic elements in FIDDLE**. The top panel displays the mass spectrum, while the bottom panel shows SHAP values indicating how each *m/z* peak contributes to the prediction of different atomic elements (C, O, N, H, P, S, F, Cl, B, Br, I, Na, K), with darker blue indicating stronger positive contribution. In this case, the labeled formula is $C_{20}H_{18}O_6$ and the predicted formula is $C_{21}H_{23}O_5$.

primarily driven by its larger scale rather than superior chemical diversity or annotation quality.

#### 2.5 Supplementary Note 4 | SHAP interpretation for model explainability

To understand how the neural network makes atomic composition predictions from MS/MS spectra, we employed SHAP (SHapley Additive exPlanations) analysis [4]. A representative subset of training spectra served as background data for the SHAP DeepExplainer. For each test spectrum, SHAP values were calculated to quantify the contribution of individual *m/z* peaks to elemental predictions.

The model wrapper fixed labeled environmental parameters during interpretation while allowing spectral features to vary. SHAP values were computed for each of the 7500 *m/z* bins (0.2 Da resolution with maximum *m/z* of 1500 Da), generating attribution scores that indicate how each spectral peak influences the prediction of different atomic elements. The resulting SHAP matrix, where rows represent elements and columns represent *m/z* bins, reveals which spectral regions drive specific elemental predictions. Positive SHAP values indicate that a peak supports the presence of a particular element.

The SHAP analysis revealed distinct element-specific attribution patterns across MS/MS spectra. Carbon and oxygen predictions were influenced by peaks throughout the entire spectrum, while hydrogen showed minimal SHAP values across all regions, likely because hydrogen atoms are ubiquitous and their presence is easily inferred from other elemental compositions. While high-intensity peaks often contributed strongly to elemental predictions, the model also assigned significant importance to lower-intensity fragments, demonstrating learned fragmentation patterns beyond simple intensity weighting. For instance, in spectrum `mona_qtof_132`, the most intense fragment at *m/z* 349.07 strongly influenced carbon attribution, while lower-intensity fragments at *m/z* 173-202 contributed significantly to nitrogen predictions.

Several limitations constrain the interpretation analysis. The binned input spectrum (0.2 Da resolution) reduces mass accuracy and may obscure individual peak contributions. Additionally, SHAP values indicate elemental presence but not specific atom counts, preventing direct fragment annotation. Without expert fragment annotation and given precursor ion exclusion, direct validation of the model's chemical reasoning remains challenging. We leave these interpretation challenges for future work. Additionally, we plan to extend our approach to molecular structure prediction in future studies, which would provide more intuitive and chemically meaningful explanations by directly linking spectral features to specific structural fragments and functional groups, thereby offering clearer insights into the model's decision-making process.
